# Data-driven model discovery and model selection for noisy biological systems

**DOI:** 10.1101/2024.10.03.616508

**Authors:** Xiaojun Wu, MeiLu McDermott, Adam L. MacLean

## Abstract

Biological systems exhibit complex dynamics that differential equations can often adeptly represent. Ordinary differential equation models are widespread; until recently their construction has required extensive prior knowledge of the system. Machine learning methods offer alternative means of model construction: differential equation models can be learnt from data via model discovery using sparse identification of nonlinear dynamics (SINDy). However, SINDy struggles with realistic levels of biological noise and does not incorporate prior knowledge of the system. We propose a data-driven framework for model discovery and model selection using hybrid dynamical systems: partial models containing missing terms. Neural networks are used to approximate the unknown dynamics of a system, enabling the denoising the data while simultaneously learning the latent dynamics. Simulations from the fitted neural network are then used to infer models using SINDy. We show, via model selection, that model discovery in SINDy with hybrid dynamical systems outperforms alternative approaches. We find it possible to infer models correctly up to high levels of biological noise of different types. We demonstrate the potential to learn models from sparse, noisy data in application to a canonical cell state transition using data derived from single-cell transcriptomics. Overall, this approach provides a practical framework for model discovery in biology in cases where data are noisy and sparse, of particular utility when the underlying biological mechanisms are partially but incompletely known.

## Introduction

Mathematical models wielded skillfully can offer great insight into biological systems. The process of constructing models, however, is typically manual and labor-intensive, and proceeds in an unscripted manner that often veers into whimsy. Data-driven approaches offer an attractive alternative: means by which models can be learnt directly from data. However, to perform such model discovery one must overcome the idiosyncrasies that biological systems present, including appropriate consideration of the extent/type of noise present in the data, and the need to evaluate results in an unbiased way.

Noise is pervasive in biological systems. Technical and biological (intrinsic and extrinsic) sources of noise must be taken into account for the accurate quantification and simulation biological dynamics. To do so, one can model a stochastic dynamical system starting from a discrete Markov process (stochastic reaction network) or a Chemical Langevin equation [1]. While treatment of the stochastic dynamics is at times necessary, models described by ordinary differential equations (ODEs), capturing the mean behavior of the underlying dynamics can be (unreasonably) effective at describing the dynamics of biological systems across many scales. A wide suite of tools readily available for the analysis of ODE models extends their utility. Here we focus on ODE models for biological systems, taking the general form *x*^*′*^ ≡ *dx/dt* = *f* (*x*(*t*)), where *x* = *x*(*t*) is the state vector of species at time *t*. Via model discovery we seek to learn ODEs for biological models, i.e. we will treat the biological noise as external to the system dynamics.

The indefatigable growth of biological datasets in size and scope offers fertile ground for the integration of machine learning and dynamical systems approaches to learn new biology [2]. Data-driven model discovery methods offer means by which model equations can be inferred directly from data, such as via sparse identification of nonlinear dynamics (SINDy) [3]. SINDy assumes that the right-hand-side of a model (e.g. *f* (*x*(*t*)) of an ODE model) can be expressed as a product of basis functions and a coefficient matrix. The goal of SINDy is then to learn the (sparse) coefficient matrix that best fits the data via regularized regression. A basis that defines the set of possible terms in the model must be constructed prior to regression; often such bases are comprised of polynomial functions, although, as we will show below, SINDy can also discover models containing non-polynomial terms (e.g. described by Hill equations [4], commonly found in biological networks). SINDy, and its extensions [5, 6], enable model discovery from a large space of possible models; previous methods able to explore topological model spaces or model properties typically operated over smaller domains [7, 8]. SINDy has been successful in a wide variety of contexts from modeling CAR-T cell therapy [9] to epidemiology [10]. Model discovery approaches extend beyond differential equation-based models as recent works combining model discovery with discrete/agent-based modeling have shown [11, 12]. Recent works have also demonstrated the potential for model discovery on noisy data [13–15], although inferring models in the presence of noise remains challenging.

In some cases, partial knowledge of a biological system is available a priori. For example: the birth or death terms of a biological species might be known, but the interaction terms for that species might be unknown. Partial knowledge can be incorporated into the model discovery framework by splitting *f* (*x*(*t*)) into a known and an unknown part, the latter of which we will approximate by a neural network (NN). We refer to such a model as a hybrid dynamical system. This approach builds upon recent successes in the literature that marry differential equation- and NN-based approaches to create new NN models that make use of prior knowledge and differential equation-solving tools [16–18]. Several recent works demonstrate the potential to infer models using neural ODEs [19–22]. Alternatively, Rackauckas et al. [23] developed means with which to combine a hybrid dynamical system with SINDy to perform partial model discovery on the *unknown* portion of a model. In this approach, the NN within a hybrid dynamical system is learned from data via an appropriate objective function, and subsequently an ODE model is inferred using SINDy on the derivatives estimated by the fitted hybrid dynamical model [23]. This approach proved successful using limited input data — short trajectories, albeit richly sampled in time. Inferring correct models however becomes difficult as the noise grows. Thus, we seek means with which to evaluate predicted models in an unbiased way given the typical unavailability of ground truth for biological systems of interest.

In this paper we develop methods for data-driven model discovery of complex biological dynamics via hybrid dynamical systems. We show that we are able to learn models from noisy biological data using only short time spans for training, i.e. species can be far from steady state. Moreover, we show how incorporating partial knowledge into the inference framework facilitates model discovery. To do so we apply a two-step model discovery framework. First, we fit the unknown dynamics using a neural network for smoothing and interpolation. Second, we use the trained neural network as input to SINDy to learn symbolic model terms. At both steps we perform model selection to search over hyperparameter space with unbiased evaluation criteria. With application to synthetic noisy data from two canonical models (Lotka-Volterra [24, 25] and the repressilator [26]) we demonstrate the advantages of hybrid dynamical systems for model discovery with complex data. We also apply these methods to a single-cell RNA-sequencing dataset describing a cell state transition, and demonstrate that we can learn models from real data.

## Materials and methods

We introduce methods for data-driven discovery of ordinary differential equation (ODE) models that govern the behavior of a dynamical system quantified by *n* noisy observations *X* = {*X*^(1)^, …, *X*^(*n*)^} sampled on a time span [0, *T*] (Fig 1A). The goal is to learn the ODEs that describe the system via a hybrid dynamical system, i.e. one for which the model may be partially but not completely known. We tackle this in a two-step approach: first learning the derivatives of *X* via the hybrid dynamical system, and then inferring the full model from the learned derivatives using SINDy. To provide unbiased means to select a model we perform model selection over the hyperparameter space, again in two steps: first for derivative learning with neural-network based approaches and then for equation learning in SINDy. We use lowercase notation for variables (e.g. *x*) and uppercase for data (e.g. *X*). We use subscripts to denote the indices of multidimensional variables, data, or functions, e.g. *x*_*i*_, the *i*-th element of *x*, or *g*_*i*_(*x*), the *i*-th component of the function *g*(*x*). One exception is *x*_0_, which we use to define the initial conditions. We use superscripts with parentheses denote labels for data, e.g. *X*^(*i*)^ refers to the *i*-th sample with a dataset, and *X*^(true)^ refers to the data representing the ground truth.

**Fig 1.**
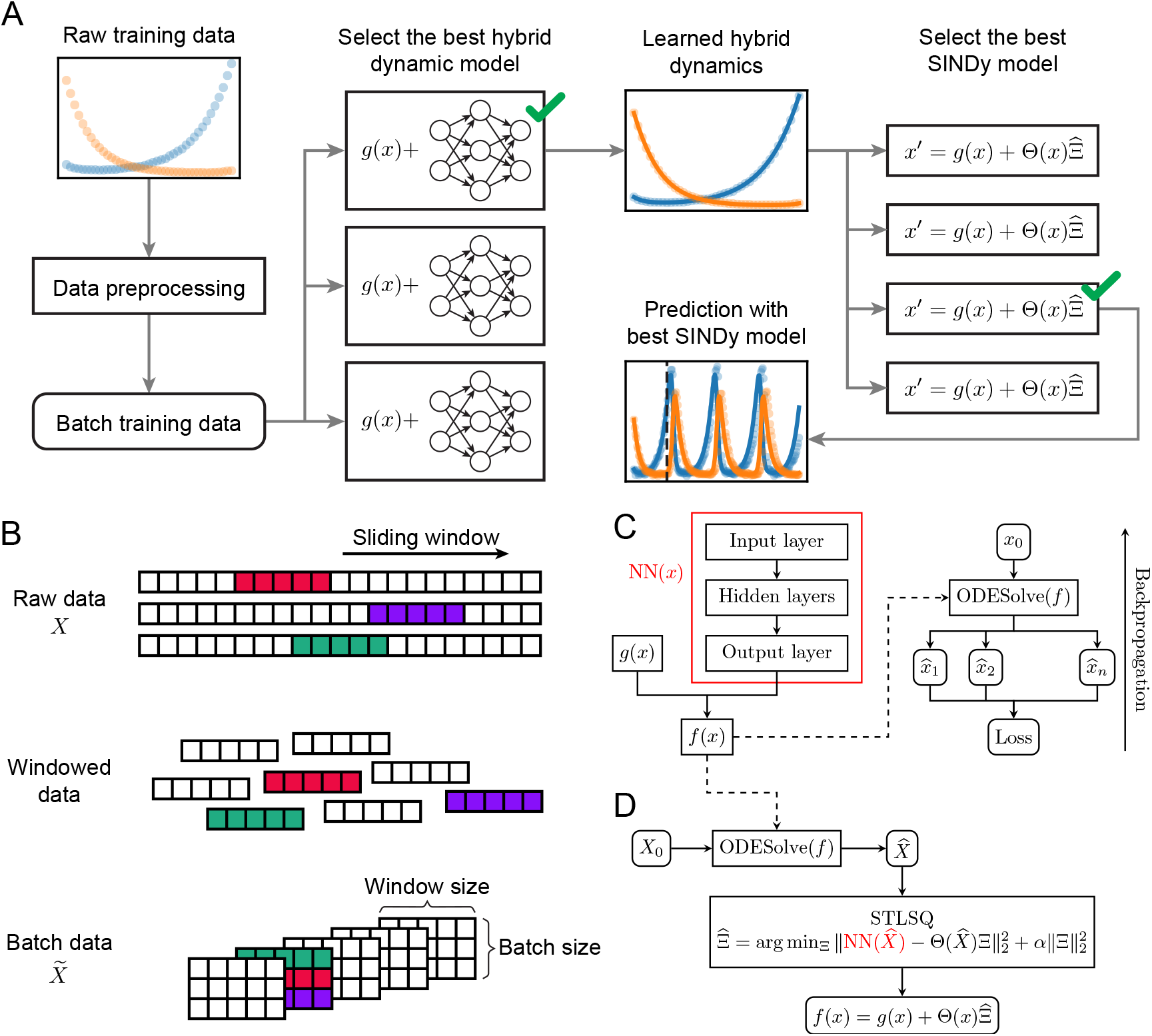
Pipeline for data-driven model discovery for dynamical systems with model selection. **A**. Overview of the pipeline. Raw data is converted to batch training data for learning hybrid dynamical models using different hyperparameters. Dynamics approximated by the best hybrid dynamical model are used for inferring ODE models with SINDy. Inferred ODE models are evaluated on model fit and extrapolation. **B**. Each sample in the raw training data is split using a sliding window to obtain samples on shorter time spans. These short samples are assembled into batches for training. **C**. A hybrid dynamical model is trained using NN(*x*) by simulating over the training time span and backpropagating, using the loss between simulated data and training data. **D**. The trained hybrid dynamical model is input to SINDy for equation-learning via the STLSQ sparse regression algorithm, and the final inferred model is output.

### Formulation of a hybrid dynamical system

A hybrid dynamical system is defined as an ODE system of the form:

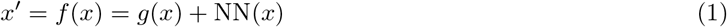

where *g*(*x*) is a closed-form function and NN(*x*) is a neural network. Note that other implementations of dynamical systems inference also use “hybrid” to describe their formulation [10, 27, 28], but these uses are distinct to that which we present here.

### Data generation

For a model given by *x*^*′*^ = *f* (*x*), we generate datasets comprised of samples for training, validation, and testing. We first simulate the model from its initial conditions, *x*_0_, (using the LSODA method from SciPy) and obtain a noise-free time series *X*^(true)^ on [0, *T*] with step size Δ*t* [29]. Noisy observations *X*^(*i*)^ are then generated according to the specified type of noise (additive or multiplicative) and the specified noise level *ϵ*. We note that we are considering sources of extrinsic (measurement) noise here, and not the intrinsic sources of noise that are also at play. In the case of additive noise, we generate observations for state variable *k* at time *t*_*j*_ from: 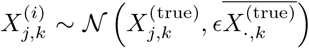, where 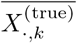 is the mean of state *k* for *X*^(true)^. That is, variation in the magnitude of the noise is constant across time points for the state *k*. In the case of multiplicative noise, we generate observations for state variable *k* from: 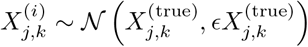. The magnitude of the noise in the observation of 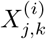 depends on the value of 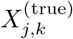 at time *t*_*j*_ only, i.e. we do not consider time-correlated noise measurements.

### Data preprocessing

We apply gradient-based optimization approaches to batches of noisy data to facilitate learning. Raw data from the training set *X* ={*X*^(1)^, …, *X*^(*n*)^} are converted into shorter time series by taking a sliding window through each *X*^(*i*)^ with a window of size *w* and a step size of 1 (Fig 1B). These short time series data are then shuffled and arranged into batches of size *b* for training (Fig 1B). We will denote the batch data as 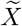.

### NN-based approaches to fit the unknown dynamics of a hybrid dynamical system

Given a hybrid dynamical system (Eq 1), following the approach taken by Rackauckas et al. in [23], we learn a neural network (NN) approximator NN(*x*) for the unknown dynamics of the system using the preprocessed training samples 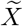 (Fig 1C). For each sample 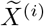 on time points {*t*_*j*_, …, *t*_*j*+*w−*1_} in a batch, we simulate *f* (*x*) from initial condition 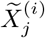,*·* to obtain the predicted dynamics 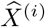 over this time range. The mean squared error (MSE) between 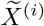 and 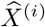 is defined as the sample loss, and the batch loss, ℒ, is defined as the mean of all the sample losses in one batch. The gradient of ℒ with respect to the weights of NN(*x*) is then computed by backpropagation.

The NN approximator is implemented in PyTorch [30]. For numerical integration we use the Dopri5 integrator as it is implemented in the torchode package [31]. For gradient descent we use the Adam algorithm to update the weights of NN(*x*) from the gradient of ℒ [32].

### Sparse regression with SINDy for model discovery from a hybrid dynamical system

We use a modified SINDy framework to infer the dynamics *f* (*x*) of an ODE system that is partially known (Fig 1D). First, given NN(*x*) trained above, we simulate the dynamics *f* (*x*) = *g*(*x*) + NN(*x*) with initial conditions taken from the training samples. By simulating over the same time span as for training, [0, *T*], with step size Δ*t* (which can provide finer temporal resolution than in training) we obtain trajectories 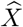 for input to SINDy. We then select a library of basis functions, Θ(*x*). The basis can contain polynomial as well as non-polynomial terms, as we will show in the examples below. SINDy then finds a sparse matrix 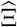 that best fits the regression problem specified by: 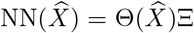. We use an implementation of the SINDy optimization algorithm Sequentially Thresholded Least Squares (STLSQ) in PySINDy that permits multi-sample input to determine 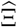. In brief, STLSQ iteratively performs ridge regression on the basis functions in Θ(*x*) with nonzero coefficients, setting coefficients to 0 if they drop below a threshold, as defined by the threshold parameter *λ* [3, 5, 6]. The full ODE model is then given by 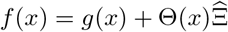.

### Model selection part I: NN-based learning

We perform two model selections steps, as we fit NN(*x*) and then 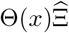 by SINDy, to allow for flexibility in the choice of hyperparameters at each step. We see in practice that one choice of hyperparameters cannot necessarily perform well on all problem types, motivating this approach. To choose the NN(*x*) that best fits the training data *X*, we allow three hyperparameters to vary: the window size (*w*), the batch size (*b*) for 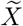, and the learning rate of the Adam algorithm (*lr*). For a predetermined set of possible values for each hyperparameter, we fit NN(*x*) on every possible combination of hyperparameter values (*w, b, lr*). For each parameter set, we train the model for 10 epochs; where in each epoch all batches in 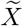 are fit. At the end of each epoch, we evaluate the current trained model using validation data. To do so, for each validation sample we simulate the trained hybrid dynamical model *f* (*x*) over the full time span to obtain the predicted dynamics; we then take observations of the predicted dynamics at the same time points as used for training. We then compute the validation loss of the sample as the MSE between the data and the predicted dynamics. The hybrid dynamical model with lowest mean validation loss is selected as the best model and retained for model discovery via SINDy at the next step.

### Model selection part II: SINDy-based learning

To find the best model discovered by SINDy, we performed sparse regression allowing for three parameters to vary: the step size Δ*t* for 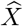, which defines the resolution of the data simulated from NN(*x*) and input to SINDy, the choice of basis function library, and the regularization parameter *α* in STLSQ. For a given model (set of ODEs recovered by SINDy) that has *k* parameters, we compute the Akaike information criterion with correction (AICc) on *n* validation samples [33]:

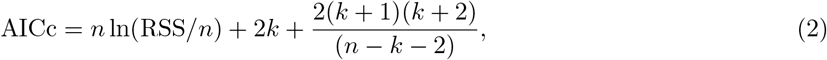

where 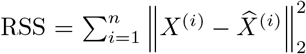 is the residual sum of squares between data *X*^(*i*)^ and model-predicted dynamics 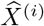. We refer to *n* ln(RSS*/n*) as the AIC term and the remainder of Eq 2 as the correction term. The model with the lowest AICc is chosen as the best model for that dataset.

### Analysis of single-cell RNA-sequencing data to infer models of the epithelial-to-mesenchymal transition

Publicly available single-cell sequencing data (read count matrices) is obtained from Cook and Vanderhyden [34] (NCBI GEO accession GSE147405) and processed in scanpy [35]. Cells with fewer than 200 genes or more than 8% mitochondrial read counts, and genes present in fewer than 3 cells are filtered out. Read counts are normalized to 10,000 and log-plus-one transformed. Batch correction is performed with ComBat [36]. Total counts, percent mitochondrial counts, and cell cycle effects are regressed out in scanpy, subsequently the data are scaled to uniform variance and highly variable genes are identified. Clustering is performed using the Leiden algorithm [37] at a resolution of 0.4. Three clusters are found at this resolution, which are identified as epithelial, intermediate, and mesenchymal cell states based on canonical gene expression. Trajectory inference is performed with diffusion pseudotime (DPT) [38]. Root nodes for trajectory inference are chosen as the cells at extreme values on the first two diffusion map components. To evaluate the uncertainty in pseudotime estimation, five independent runs of DPT are performed each with different epithelial root nodes.

The median values across the five runs are used as the pseudotime value for each cell, and the standard deviation around these values is estimated. To estimate cell state proportions during EMT, the pseudotime is divided into twelve bins. The cells of each state in each bin are counted, and the counts are converted into proportions based on the total number of cells per time point.

Datasets for inferring ODE models for cell state transition dynamics are generated from the calculated cell proportions of all states. For each observation *X*^(*i*)^ in a dataset, its value for cell state *k* at pseudotime bin *j*, denoted as 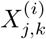, is drawn from a truncated distribution based on 𝒩(*μ*_*j,k*_, *σ*_*j,k*_), where *μ*_*j,k*_ is the mean proportion of cells in state *k* and *σ*_*j,k*_ the standard deviation. Each 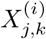 is constrained so that it is within one standard deviation of the base distribution and also within [0, 1]. Once values for two cell states are generated, the proportion of the remaining state is computed by subtracting the generated values from 1.

## Results

### A model selection framework for robust model discovery with hybrid dynamical systems

We will perform model discovery on dynamical models of the form *x*^*′*^ = *f* (*x*) with *x* ∈ ℝ^*d*^, where a partial but incomplete closed-form expression of *f* (*x*) is given. This can be represented as a hybrid dynamical system (Eq 1):

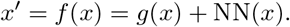

Here, *g* : ℝ^*d*^ → ℝ^*d*^ is a function known in closed-form, and NN : ℝ^*d*^ → ℝ^*d*^ is a neural network that models the unknown latent dynamics, which will be learnt from data and then input to SINDy (Fig 1A). In the case that we have no prior knowledge about the model (no terms in *f* (*x*) are known), we set *g*(*x*) = 0 and learn the entire right hand side via NN(*x*). From NN(*x*), we then run SINDy, i.e. perform regression on NN(*x*) = Θ(*x*)Ξ to find a sparse coefficient matrix 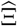 over the basis Θ(*x*) that characterizes the learnt model (Fig 1D). The full ODE model is then given by 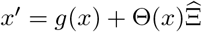.

We perform model selection at both the NN and the SINDy steps to select the model that best describes the data. To demonstrate the pipeline, we will use the Lotka-Volterra model as an example (Fig 2A), which is given by [24, 25]:

**Fig 2.**
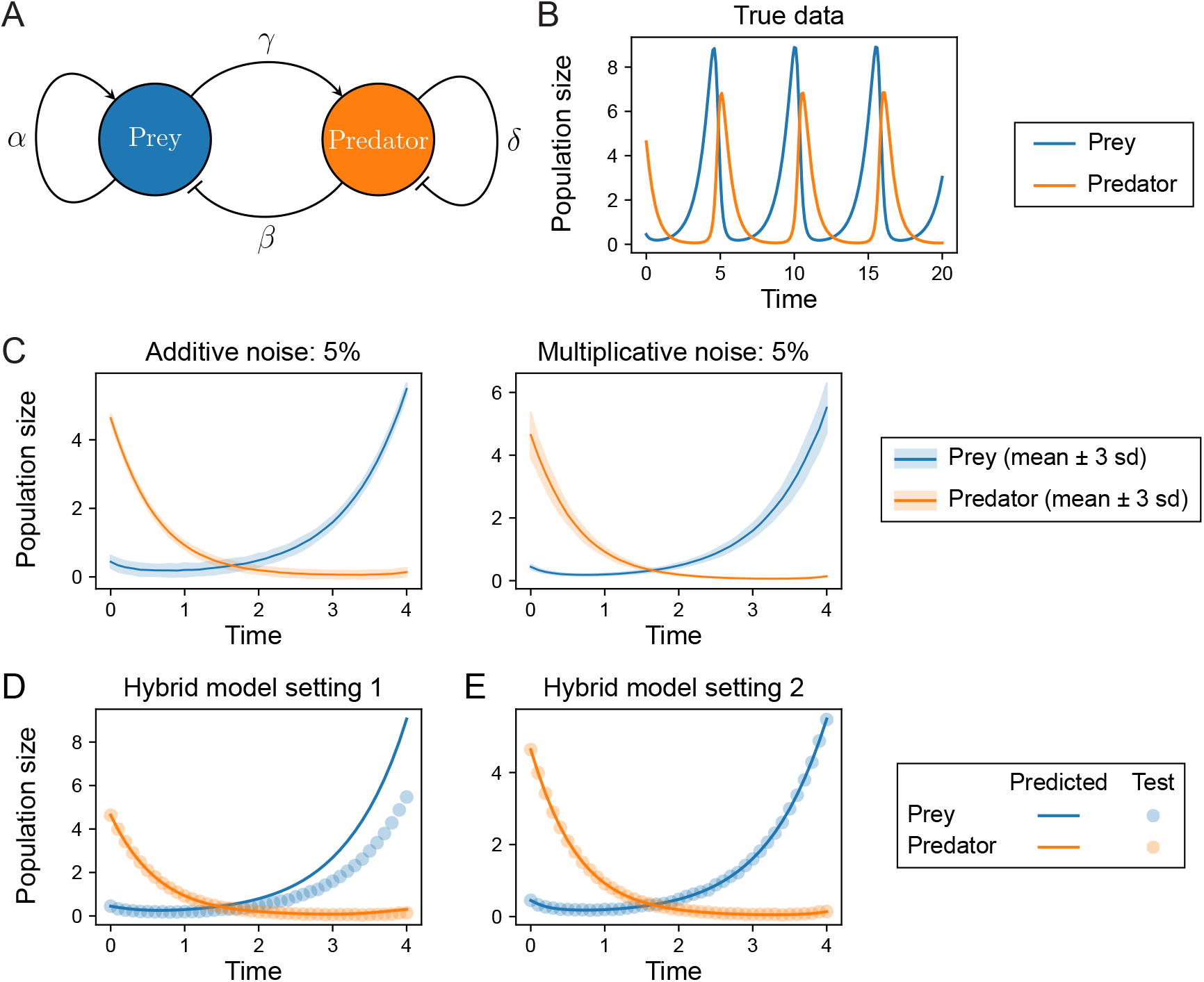
Evaluation of hybrid dynamical model fits on the Lotka-Volterra model. **A**. The Lotka-Volterra system describes predator-prey relationships in an ecosystem, here parameterized by (*α, β, γ, δ*). **B**. Example simulation with parameters (1.3, 0.9, 0.8, 1.8). The population dynamics oscillate at a stable limit cycle. **C–D**. To evaluate model discovery methods, additive or multiplicative noise is added to the underlying deterministic dynamics at different noise levels. The mean trajectories of 200 samples for each noise model are shown; ribbons represent ±3 s.d. **E**. Comparison of fits using hybrid dynamical models. In setting 1, good fits to the data were not obtained (high validation loss). Training parameters: learning rate 0.001; window size 10; batch size 10. In setting 2, a good fit to the data was obtained. Training parameters: learning rate 0.01; window size 5; batch size 5.

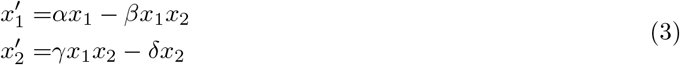

In Eqs 3, *x*_1_ is the population size of the prey and *x*_2_ is the population size of the predators.

Our model selection strategy consists of two stages. In the first, we determine the best NN(*x*) in Eq 1 to model the latent dynamics. For the Lotka-Volterra model, we assumed that the growth/death terms were known, and sought to learn the interaction terms via NN : ℝ^2^ → ℝ^2^ with:

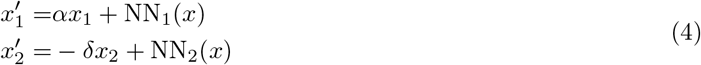

The neural network NN(*x*) in Eqs 4 had an input layer of length 2 and an output layer of length 2. Between those two layers, we added one fully-connected hidden linear layer of 8 neurons, omitting greater depth to avoid overfitting due to model complexity. For each dataset, we trained NN(*x*) with different learning rates on samples processed by various window and batch sizes (see Material and methods); for the Lotka-Volterra model the hyperparameter values used in model selection are given in Table 1.

**Table 1.**
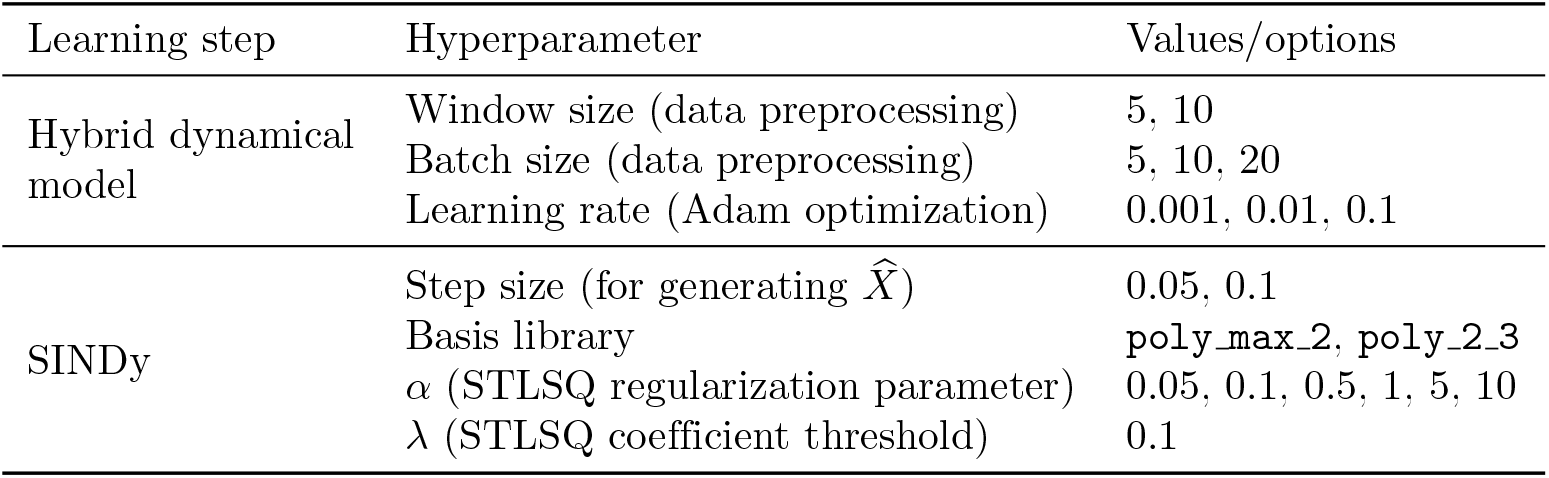
Model selection configuration for Lotka-Volterra.

In the second stage of our model selection approach, we sought to discovery accurate ODE models by SINDy. We evaluated models by the Akaike information criterion with correction (AICc, Eq 2), which estimates the model prediction error with a penalty term to account for the number of model parameters [33]. For the Lotka-Volterra model, we considered three hyperparameters for model selection. The first hyperparameter was the step size used in the generation of training data 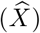 for SINDy from the fitted hybrid dynamical model. We note that generating trajectories with an arbitrary step sizes (interpolation) is only possible because of the fitted hybrid dynamical system; methods such as that of finite differences only compute derivatives on the time points provided. The second hyperparameter was the library of basis functions. Here, we assumed that the possible interaction terms were all polynomial, but with unknown order. We constructed two basis libraries consisting of polynomial terms of different orders: the first included all polynomial terms up to order 2 (denoted `poly_max_2`); the second included only polynomial terms of orders 2 and 3 only (denoted `poly_2_3`), which was motivated by the observation that we sought to infer interaction terms, i.e. the first-order terms in the model may be already completely known. The third hyperparameter was the regularization parameter *α* for STLSQ. Table 1 gives the full details of the hyperparameter space explored for model selection. We kept *λ*, the STLSQ coefficient threshold, constant at *λ* = 0.1, i.e. only basis terms with coefficients greater than 0.1 are retained in the final model.

### Lotka-Volterra models can be inferred from noisy data

To test whether we could correctly infer the Lotka-Volterra model (Fig 2A) from noisy data, we generated datasets from a single set of true parameters, {*α, β, γ, δ*} = {1.3, 0.9, 0.8, 1.8} and initial conditions *x*_0_ = [0.4425, 4.6281]. Under these conditions the system oscillates, reaching a stable limit cycle with a period of approximately five time steps (Fig 2B). A total of 12 datasets were generated by considering either additive or multiplicative noise at six different noise levels: 0.1%, 0.5%, 1%, 5%, 10%, and 20% (examples for the 5% noise level are shown in Fig 2C–D). Each dataset comprised 200 training samples and 50 validation samples on *t* = [0, 4], and 50 test samples on *t* = [0, 20], all generated with step size Δ*t* = 0.1. This Lotka-Volterra example model was studied by Rackauckas et al. [23] to demonstrate how a hybrid formulation could be used to approximate derivatives for use with SINDy. Here we sought to learn from noisy biological datasets that contain multiple measurements of each species by learning the dynamics from all the data together, rather than fitting a model to a single trajectory. We also implemented a model selection strategy in order to find models that generalize on both seen and unseen data.

We found wide variation in the results of training the hybrid model (Eqs. 4) based on the choice of hyperparameters. For example, at 1% additive noise, a learning rate of 0.001 produced underfit models with high validation loss (Fig 2E), whereas higher learning rates quickly attained low validation losses and fit the data well (Fig 2F). We examined hyperparameter values used to attain lowest validation loss for all datasets and found that the best hyperparameter combination varied by datasets (S1 Table), highlighting the utility of a flexible approach. Overall, we found this approach to improve the fits of hybrid dynamical models in an automated manner; avoiding the need for ad hoc approaches in hyperparameter choice for NN(*x*).

Given the best fit NN model, we simulated training data and performed SINDy for model selection for all configurations specified in Table 1, to produce a total of 24 discovered models. We considered a model correct if it had the correct topology, i.e. correct terms in Eqs 3 with no extra terms; the parameter values did not have to be accurately inferred, although we saw in practice that the inferred parameters of most models with correct terms were close to the true values. For datasets of both additive and multiplicative noises up to 5% noise level, the ODE models with lowest AICc were all correct (Table 2). Those models could be used to accurately predict future states for test data on *t* = [0, 20] (Fig 3A–B, D–E). For 10% and 20% noise levels, extra terms were recovered in models with lowest AICc (Table 2); those models failed to extrapolate well beyond *t* = 4 (Fig 3C, F), exhibiting damped oscillations. Nonetheless, correct models could still be successfully recovered at 10% noise level (S2 Table). Those models produced stable oscillations beyond *t* = 4, albeit with small inaccuracies in the period and amplitude (Fig 3G–H). In several cases, we found that the best-fit models were inferred from a hybrid dynamical model simulated at finer temporal resolution (step size Δ*t* = 0.05) than the data, highlighting that the hybrid dynamical model approach allowed us to obtain better results as it permits data interpolation with arbitrary step size (S3 Table).

**Table 2.** Best ODE model inferred from Lotka-Volterra data at each noise level.

| Noise type | Noise level | Inferred model |
| --- | --- | --- |
| Additive | 0.1% | $x'_1 = 1.3x_1 - 0.907x_1x_2$<br>$x'_2 = -1.8x_2 + 0.804x_1x_2$ |
| Additive | 0.5% | $x'_1 = 1.3x_1 - 0.911x_1x_2$<br>$x'_2 = -1.8x_2 + 0.780x_1x_2$ |
| Additive | 1% | $x'_1 = 1.3x_1 - 0.906x_1x_2$<br>$x'_2 = -1.8x_2 + 0.781x_1x_2$ |
| Additive | 5% | $x'_1 = 1.3x_1 - 0.893x_1x_2$<br>$x'_2 = -1.8x_2 + 0.814x_1x_2$ |
| Additive | 10% | $x'_1 = 1.3x_1 + 0.206x_2^2 - 0.842x_1x_2 - 1.044x_1x_2^2$<br>$x'_2 = -1.8x_2 + 0.924x_1x_2$ |
| Additive | 20% | $x'_1 = 1.3x_1 + 0.199x_2^2 - 0.525x_1^2x_2 - 1.707x_1x_2^2$<br>$x'_2 = -1.8x_2 + 0.798x_1x_2 + 0.237x_1x_2^2$ |
| Multiplicative | 0.1% | $x'_1 = 1.3x_1 - 0.889x_1x_2$<br>$x'_2 = -1.8x_2 + 0.826x_1x_2$ |
| Multiplicative | 0.5% | $x'_1 = 1.3x_1 - 0.910x_1x_2$<br>$x'_2 = -1.8x_2 + 0.761x_1x_2$ |
| Multiplicative | 1% | $x'_1 = 1.3x_1 - 0.935x_1x_2$<br>$x'_2 = -1.8x_2 + 0.814x_1x_2$ |
| Multiplicative | 5% | $x'_1 = 1.3x_1 - 0.836x_1x_2$<br>$x'_2 = -1.8x_2 + 0.846x_1x_2$ |
| Multiplicative | 10% | $x'_1 = 1.3x_1 - 0.884x_1x_2 - 0.333x_1^2x_2$<br>$x'_2 = -1.8x_2 + 0.883x_1x_2$ |
| Multiplicative | 20% | $x'_1 = 1.3x_1 + 0.127x_1 - 1.194x_1x_2$<br>$x'_2 = -1.8x_2 + 0.502x_2 + 0.247x_1x_2 - 0.198x_2^2$ |

Terms colored in green are known terms in Eqs 4.

**Fig 3.**
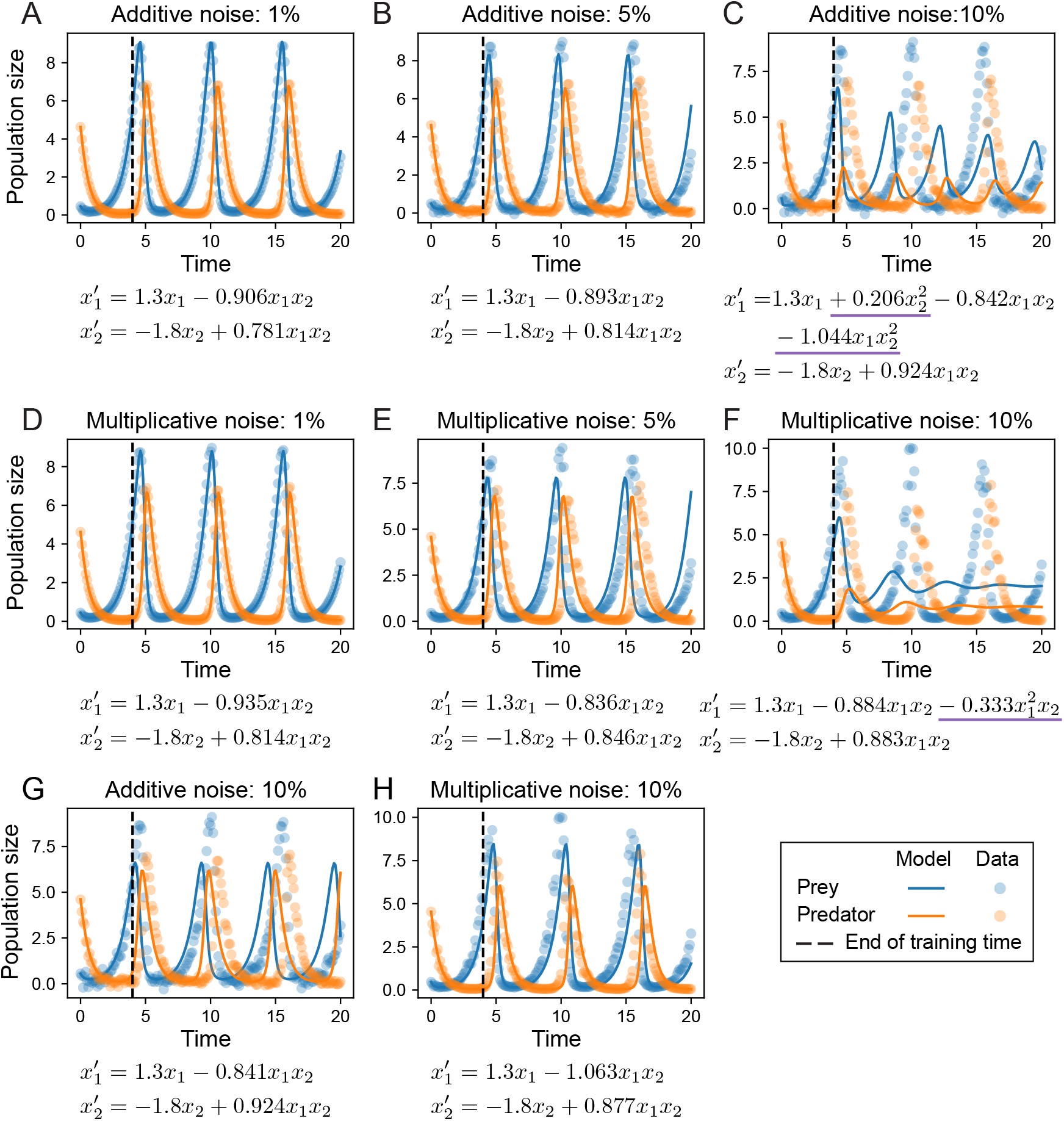
Inferred models from Lotka-Volterra datasets. For each model, the first term in 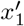 and 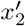 is known and the remaining terms are inferred. Underline denotes terms that are incorrect relative to the true model (Eqs 3). **A–C**. Inferred models with lowest AICc for datasets with additive noise, at 1, 5 and 10%. **D–F**. Inferred models with lowest AICc for datasets with multiplicative noise at at 1, 5 and 10%. **G**. The model with correct terms could be inferred at 10% additive noise, but was not ranked highest by AICc. **H**. The model with correct terms could be inferred at 10% multiplicative noise, but was not ranked highest by AICc.

### Incorporating prior knowledge for model discovery improves SINDy learning

To assess the performance of the hybrid dynamical system formulation for model discovery (which incorporates partial prior knowledge of the system) we compared it to two alternative methods, each of which do not incorporate prior system knowledge. The two alternative methods tested were: SINDy run in a single step on the entire data (referred to as “base SINDy”). This takes as input the training data *X* and estimates the derivatives *X*^*′*^ by finite differences [5, 6]. The second method was a “pure NN” approach, in which the entire derivative was approximated by a NN that is then input to SINDy, i.e. *x*^*′*^ = *f* (*x*) = NN(*x*), where we use the same model selection configuration as specified in Table 1. For fair comparison, we also perform a model selection step for base SINDy over different choices of basis and STLSQ regularization parameters; we could not search over different step sizes as base SINDy does not interpolate.

We compared the best correct models that were recovered by each method via the AICc. Base SINDy inferred correct models at all levels of additive noise tested, but could not find correct models for multiplicative noise *>* 1% (Table 3, S4 Table). Of note, base SINDy was the only method able to discover the correct model for high levels (20%) of additive noise. The pure NN approach inferred correct models for most levels of additive noise (0.1%, 1%, 5%, 10%) and for low levels of multiplicative (up to 1%) (Table 3, S5 Table). The hybrid formulation achieved the best performance overall, inferring correct models for 10 datasets out of 12, at all noise levels except for 20% (additive or multiplicative; Table 3). The hybrid formulation was the only able to handle higher levels of multiplicative noise, successfully discovering correct models at 5% and 10%. In terms of the quality of models discovered, the hybrid formulation had the lowest AICs for 7/12 datasets and was within a small margin of error for an additional two datasets, confirming that overall the hybrid formulation outperformed base SINDy or pure NN approaches to infer models from noisy data (Table 3).

**Table 3.** Lowest AICc values of models inferred from Lotka-Volterra datasets using various methods. Hybrid refers to the hybrid dynamical model formulation (Eqs 4). Bold denotes the lowest AICc at a given noise type/level. Model AICc is n/a if a fit was not found at that noise level

| Noise type | Noise level | Base SINDy | Pure NN | Hybrid |
| --- | --- | --- | --- | --- |
| Additive | 0.1% | -309 | **-433** | -369 |
| Additive | 0.5% | -299 | N/A | **-365** |
| Additive | 1% | -283 | -300 | **-368** |
| Additive | 5% | -178 | -39 | **-216** |
| Additive | 10% | -105 | **-147** | -146 |
| Additive | 20% | **-26** | n/a | n/a |
| Multiplicative | 0.1% | -298 | -332 | **-372** |
| Multiplicative | 0.5% | -254 | -318 | **-393** |
| Multiplicative | 1% | **-215** | -81 | -208 |
| Multiplicative | 5% | n/a | n/a | **-137** |
| Multiplicative | 10% | n/a | n/a | **-59** |
| Multiplicative | 20% | n/a | n/a | n/a |

To analyze the relative performance of each method in more depth, we studied the derivatives that were inferred by each method, as these are crucial to SINDy. We found that the derivatives for both base SINDy and a pure NN approach deviated from the true values, especially for multiplicative noise (Fig 4A–B). The derivatives inferred by our hybrid approach (where we infer a partial model) were closer to the true values overall (Fig 4C), with only small deviations for multiplicative noise near the end of the time window. This goes to explain in part how the hybrid approach is able to infer more accurate models overall in Table 3.

**Fig 4.**
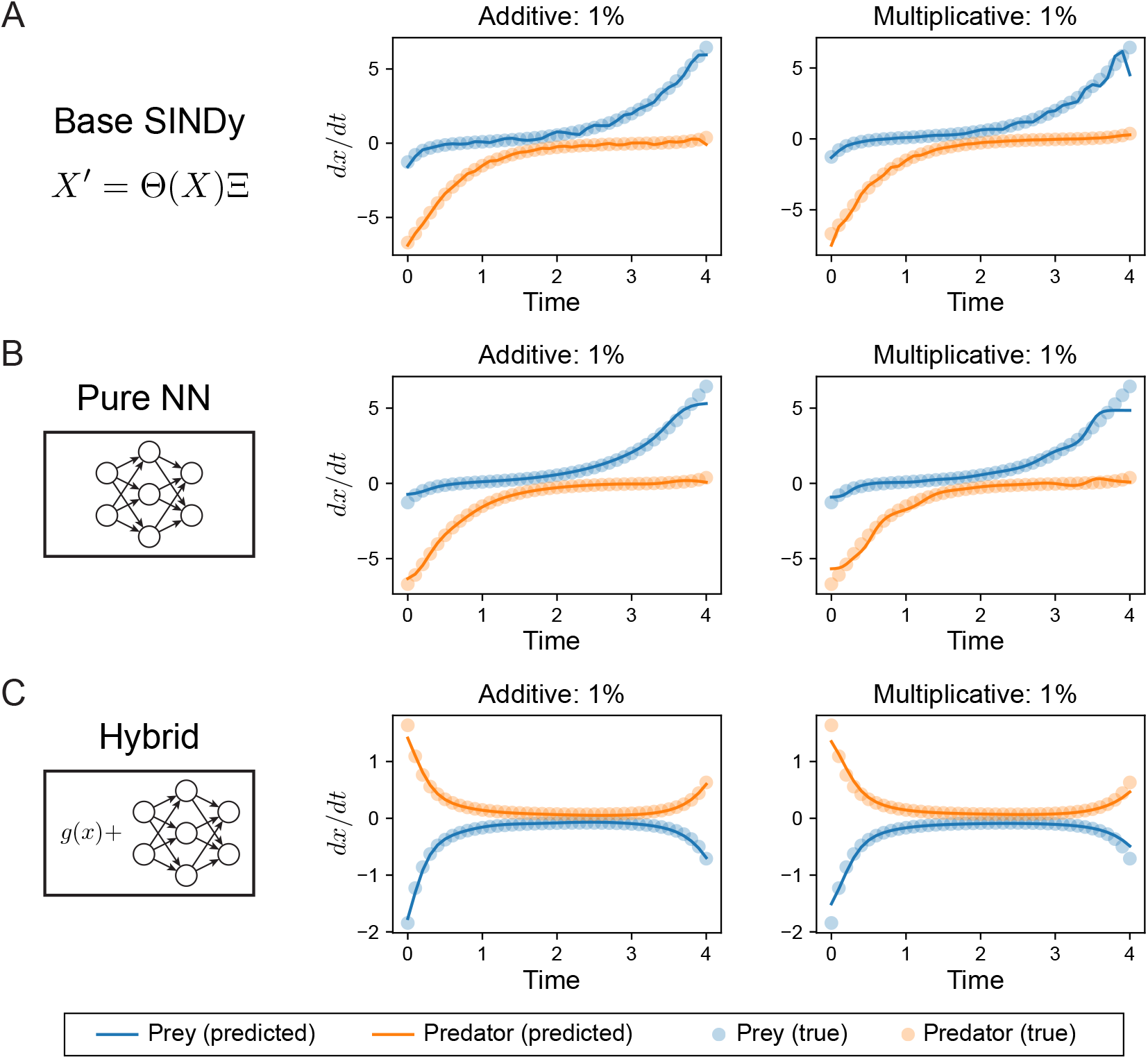
Comparison of derivatives approximated from Lotka-Volterra datasets using various methods. **A**. Derivatives approximated by base SINDy, which uses the method of finite differences. **B**. Derivatives approximated by a pure neural network formulation, in which the full right-hand side of an ODE system is approximated by a neural network. **C**. Derivatives approximated by a neural network in the hybrid model; where the neural network is fitted to partial dynamics (Eqs 4).

### Model discovery from noisy data for biological reaction networks with complex dynamics

We next turn our attention to a larger model with complex dynamics, as a more challenging test of model discovery with hybrid dynamical systems. The repressilator [26] describes a synthetic transcriptional network that can produce stable oscillations. Originally constructed in E. coli, it consists of three proteins that each repress their neighbor, thus coupling all three in a negative feedback loop (Fig 5A). It can be modeled with six species: the mRNA and protein products of each gene, or one simplify the model to consider one variable per species. Taking the latter approach, the system can be described by the following minimal model:

**Fig 5.**
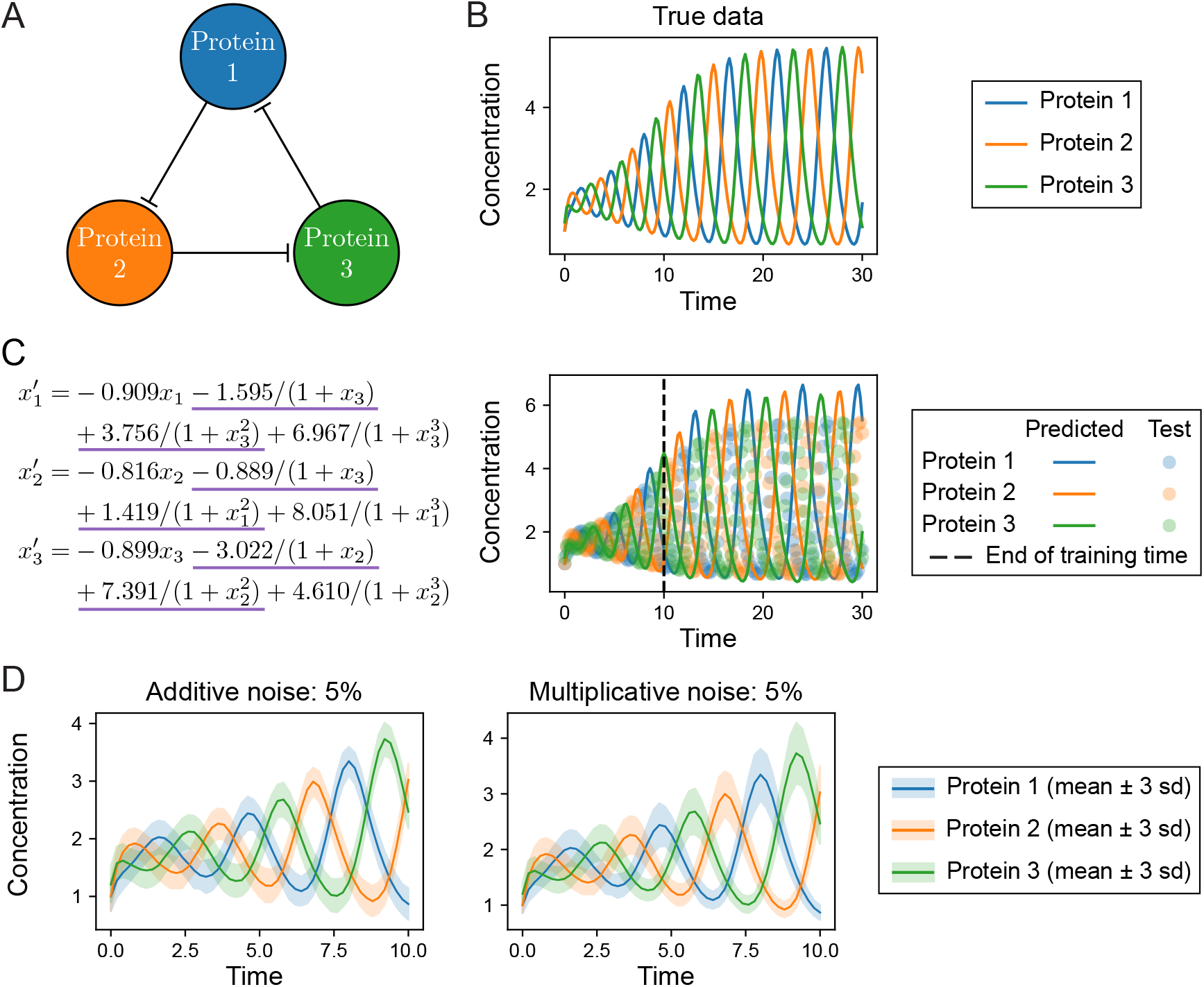
Evaluation of base SINDy fits and data generation for the repressilator model. **A**. The repressilator models describes a coupled negative feedback loop between three interacting proteins. **B**. Example simulation of the repressilator model without noise. The concentrations of each of the three proteins oscillate and the system eventually reaches a stable limit cycle. **C**. Evaluation of base SINDy to infer repressilator models from noise-free data on *t* = [0, 10]. Inferred equations (left) and simulation of the inferred model on *t* = [0, 30] (right). Underlined terms are incorrect in comparison to the true repressilator model (Eqs 5). **D–E**. To evaluate model discovery methods, additive or multiplicative noise is added to the underlying deterministic dynamics at different noise levels. The mean trajectories of 200 samples for each noise model are shown; ribbons represent ±3 s.d.

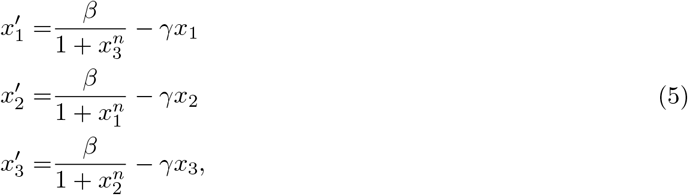

where *β* and *γ* represent the basal production and degradation rates, respectively, and where we have assumed symmetry between the species. Below, we will set the degradation rate for each protein as *γ* = 1 and assume these are known for each species. The remaining terms in Eqs 5 represent the inhibitions of each protein by its neighbor in the network. The inhibition terms are represented by Hill functions of order *n*, where we have assumed a half maximal effective concentration of each protein *k* = 1. We studied a specific parameterization of the repressilator system with *n* = 3, *β* = 10 and initial conditions *x*_0_ = [1, 1, 1.2]. At this point, the system displays oscillations that increase in amplitude for the first 15 time steps (units are dimensionless) and then reaches a stable limit cycle thereafter (Fig 5B). We tested whether we would discover correct models for the repressilator (Eqs 5) from data on *t* = [0, 10], a time interval that does not let the system reach its steady state.

We first tested whether we could discover correct models for the repressilator using a direct SINDy approach (base SINDy). To construct a basis for regression in SINDy, we made the assumptions that: the degradation term for each protein is linear; proteins interact only via inhibition; and that the strength of each inhibitory interaction can be characterized by a Hill function 1*/*(1 + *u*^*n*^) of order *n*, with *n* ∈ {1, 2, 3}. I.e. we seek to discover the interaction structure of the network for the pairs of proteins: who interacts with whom, and in what direction. Given these assumptions, the basis library for SINDy consisted of the Hill functions {1*/*(1 + *x*_*i*_), 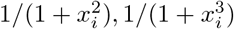} and the linear functions {*x*_*i*_} for *i* = 1, 2, 3; a total of 12 basis functions and 36 coefficients to be determined. In testing model discovery with SINDy on this basis, we realized that it was difficult to set *λ*, the basis coefficient threshold for inclusion of that term. For *λ* close to or greater than 1, the (correct) linear terms would either be eliminated or have coefficients much larger than the true values. For *λ* ≪ 1, all of the discovered models would admit extra Hill terms with wrong orders and small coefficients. For example, with *λ* = 0.5 and true data generated noise-free and sampled every 0.2 seconds in *t* = [0, 10], the discovered model is shown in Fig 5C. This model has Hill terms of orders *n* = 1 and *n* = 2 and cannot accurately predict future states (Fig 5C). This highlights the challenge in estimating derivatives for SINDy, in this case even before any noise is added to the data.

The hybrid formulation overcomes the challenge of threshold choice in SINDy. We separate the linear (known) degradation terms from the unknown interaction terms in the hybrid dynamical system with NN : ℒ^3^ → ℒ^3^ to give:

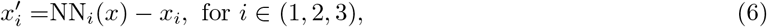

where NN(*x*) is a neural network with two fully-connected hidden linear layers of 8 neurons each. In the hybrid system, the choice of *λ* no longer affects the linear terms since these are known; and need to learn only the interaction terms from the latent dynamics approximated by NN(*x*). To do so we used a basis consisting of inhibitory terms, i.e. Hill functions between pairs of proteins with different Hill coefficients:

{1*/*(1 + *x*_*i*_), 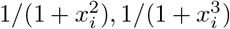}. In addition, our model selection scheme for SINDy would recover models using basis libraries of Hill functions of the same order. We will use hill n for a basis library of Hill functions of order *n* only and hill max n for a basis library of all Hill functions up to order *n*. The full configuration for model selection on repressilator models is given in Table 4.

**Table 4.** Model selection configuration for the repressilator model.

| Learning step | Hyperparameter | Values/options |
| --- | --- | --- |
| Hybrid dynamical model | Window size (data preprocessing) | 5, 10 |
|  | Batch size (data preprocessing) | 5, 10, 20 |
|  | Learning rate (Adam optimization) | 0.001, 0.01, 0.1 |
| SINDy | Step size (for generating $\hat{X}$) | 0.1, 0.2 |
|  | Basis library | hill_1, hill_2, hill_3, hill_max_3 |
| | $\alpha$ (STLSQ regularization parameter) | 0.05, 0.1, 0.5, 1.0, 5.0, 10.0 |
| | $\lambda$ (STLSQ coefficient threshold) | 1.0 |

To test whether we can successfully discover the repressilator model from noisy data, we again generated datasets with additive or multiplicative noise at six different noise levels from 0.1 – 20% (example for 5% noise shown in Fig 5D–E). Each dataset had 200 training samples and 50 validation samples on *t* = [0, 10], as well as 50 test samples on *t* = [0, 30], all sampled with step size Δ*t* = 0.2. As for the example with the Lotka-Volterra model, we considered an inferred ODE model correct if it identified all terms of Eqs 5 with no extra terms (i.e. correct topology). We did require that “correct models” had highly accurate parameter values, although we found in practice that they often were very close to the true values.

We found that correct models were inferred for all noise types and levels up to 1%, and for multiplicative noise up to 5% (Table 5). The inferred models could accurately predict future states (Fig 6A–B, D–E). For higher noise values, the inferred models with lowest AICc did not correctly recover Eqs 5 and could not predict future dynamics beyond the training time span (Fig 6C, F). However, analysis of the top 10 inferred models for each noise type/level at higher levels of noise (S6–S10 Tables) revealed that in many cases correct models could be found. For additive noise at 5% and 10% and for multiplicative noise at 10%, correct ODE models were discovered in the top 10 predicted models, e.g. Models 4 and 5 in S6 Table, Models 6 and 7 in S7 Table, Models 7 and 8 in S8 Table. Simulations of these models showed that — despite larger parameter variation from the true values — they produced stable limit cycles with oscillations close to the true data well beyond the time range used for training (Fig 6G–H).

**Table 5.**
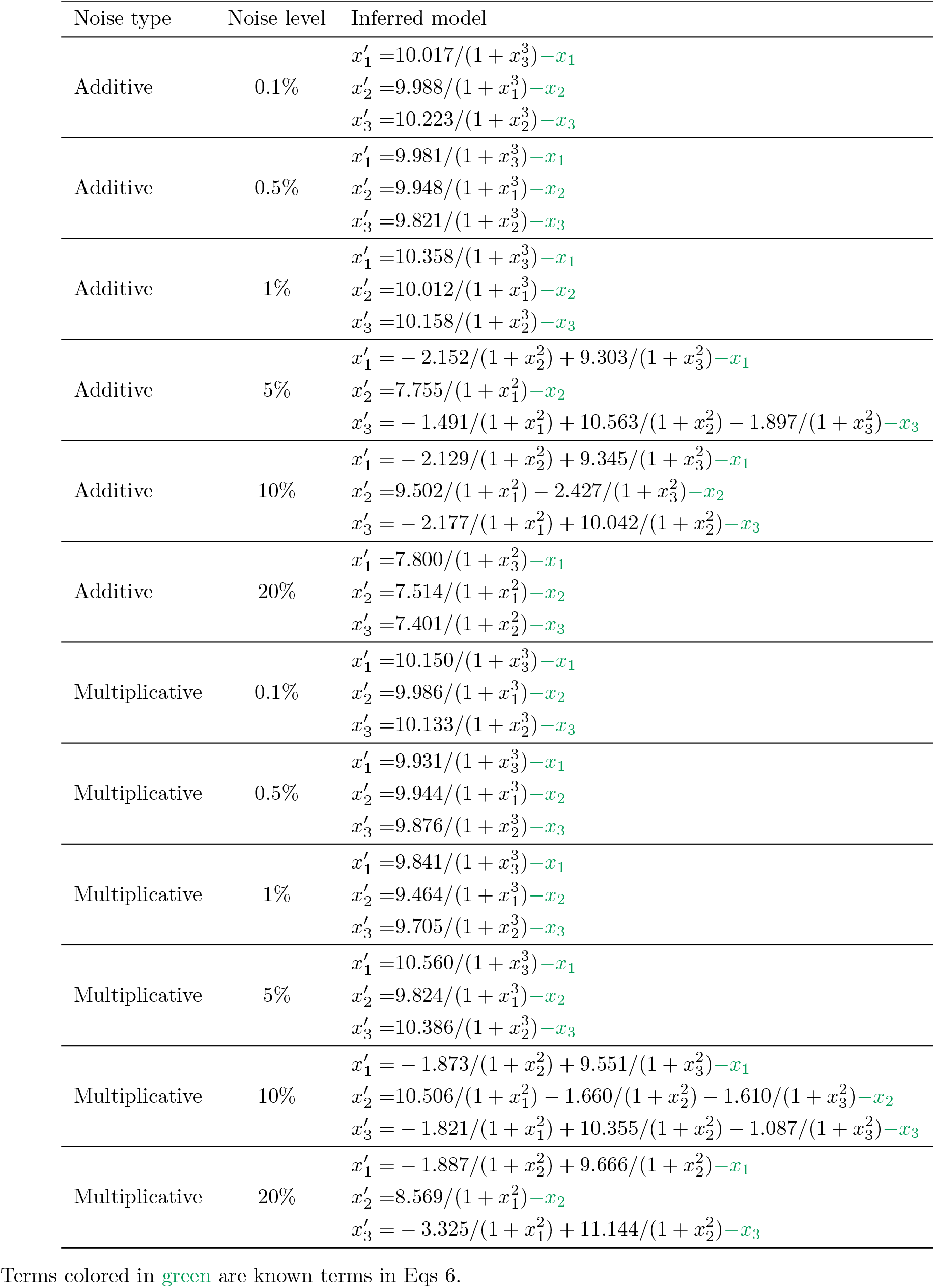
Best ODE model inferred from repressilator data at each noise level.

**Fig 6.**
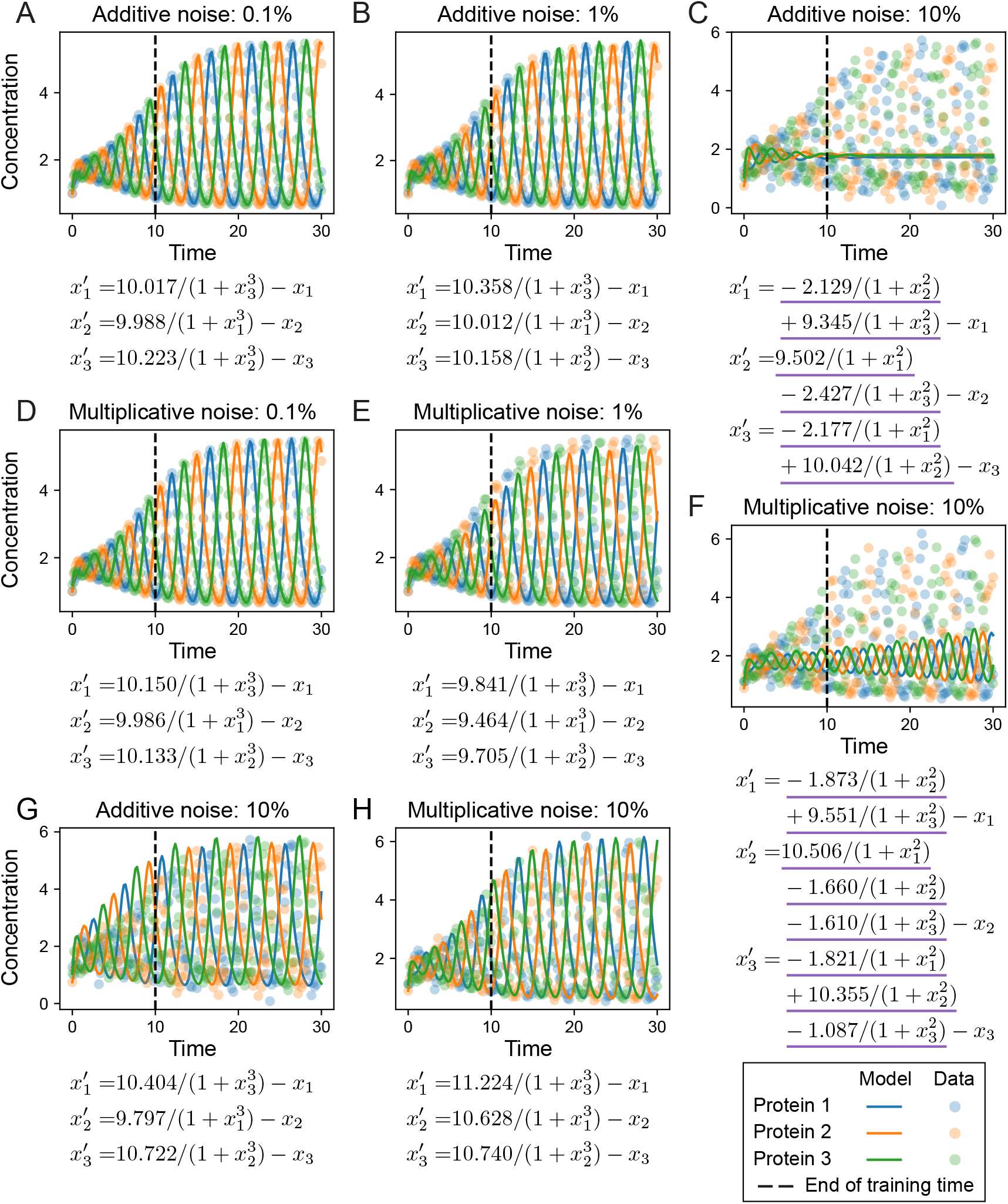
Inferred models inferred from repressilator datasets. For each model, the linear decay terms are known and the remaining terms are inferred. Underline denotes terms that are incorrect relative to the true model (Eqs 5). **A–C**. Inferred models with lowest AICc for datasets with additive noise, at 0.1, 1 and 10%. **D–F**. Inferred models with lowest AICc for datasets with multiplicative noise at at 0.1, 1 and 10%. **G**. The model with correct terms could be inferred at 10% additive noise, but was not ranked highest by AICc. **H**. The model with correct terms could be inferred at 10% multiplicative noise, but was not ranked highest by AICc.

At the 20% noise level, there were no correct models in the top 10 predicted for either additive or multiplicative noise. However, even at this high noise level, inferred models with low AICc scores contained aspects of the true system. Some inferred models contained Hill terms of the correct order (*n* = 3) but with an extra term (e.g. Models 4 and 5 in S9 Table, or Models 7 and 8 in S10 Table). In these models the correct terms had coefficients close to true values (*β* = 10) and the incorrect terms had coefficients small in magnitude. Additionally, models were predicted that contained Hill terms of the wrong order but which recapitulated the correct repressilator network. As an example, Model 1 (S9 Table) inferred the correct network topology:

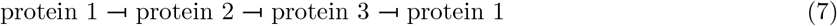

although with lower cooperativity (smaller Hill term: *n* = 2) than the true system. We saw again in the case of models with incorrect Hill terms, that if they contained additional (incorrect) network edges (e.g. Model 1 in S10 Table), these would likely have much smaller coefficients than the terms that represented correct edges in the network (Eq 7).

This analysis highlights useful applications and limitations of SINDy to recover complex models from noisy data. At high noise levels it might not be possible to infer correct models with confidence. Nonetheless, through careful analysis of the predicted models in light of: model topology, parameter values (coefficients), and the consistency of ranked model predictions (e.g. if models show the same network but with different Hill coefficients) we can gain predictions regarding model structure and interactions for further testing.

### Model discovery of cell state transition dynamics from single-cell transcriptomics

To demonstrate the potential of model discovery to infer the structure of biological systems from real data, we applied the hybrid dynamical system approach to a dataset characterizing the epithelial-to-mesenchymal transition (EMT), a canonical cell state transition of importance in development, wound healing, and cancer [39–41]. We studied EMT induced by TGF-*β* in lung carcinoma cells (A549). Theoretical and experimental evidence has indicated that intermediate states exist along the EMT spectrum [42, 43]. We found evidence for one stable intermediate cell state in this dataset (see Methods) and sought to infer models describing the dynamics of three cell populations over pseudotime: epithelial (E), intermediate (I), and mesenchymal (M) (Fig 7A). A direct transition from *E* → *M* might also occur, as well as reverse transitions. By projecting the cell states over pseudotime (see Methods) we obtain the cell population pseudo-dynamics, from which we can infer models of EMT (Fig 7B).

**Fig 7.**
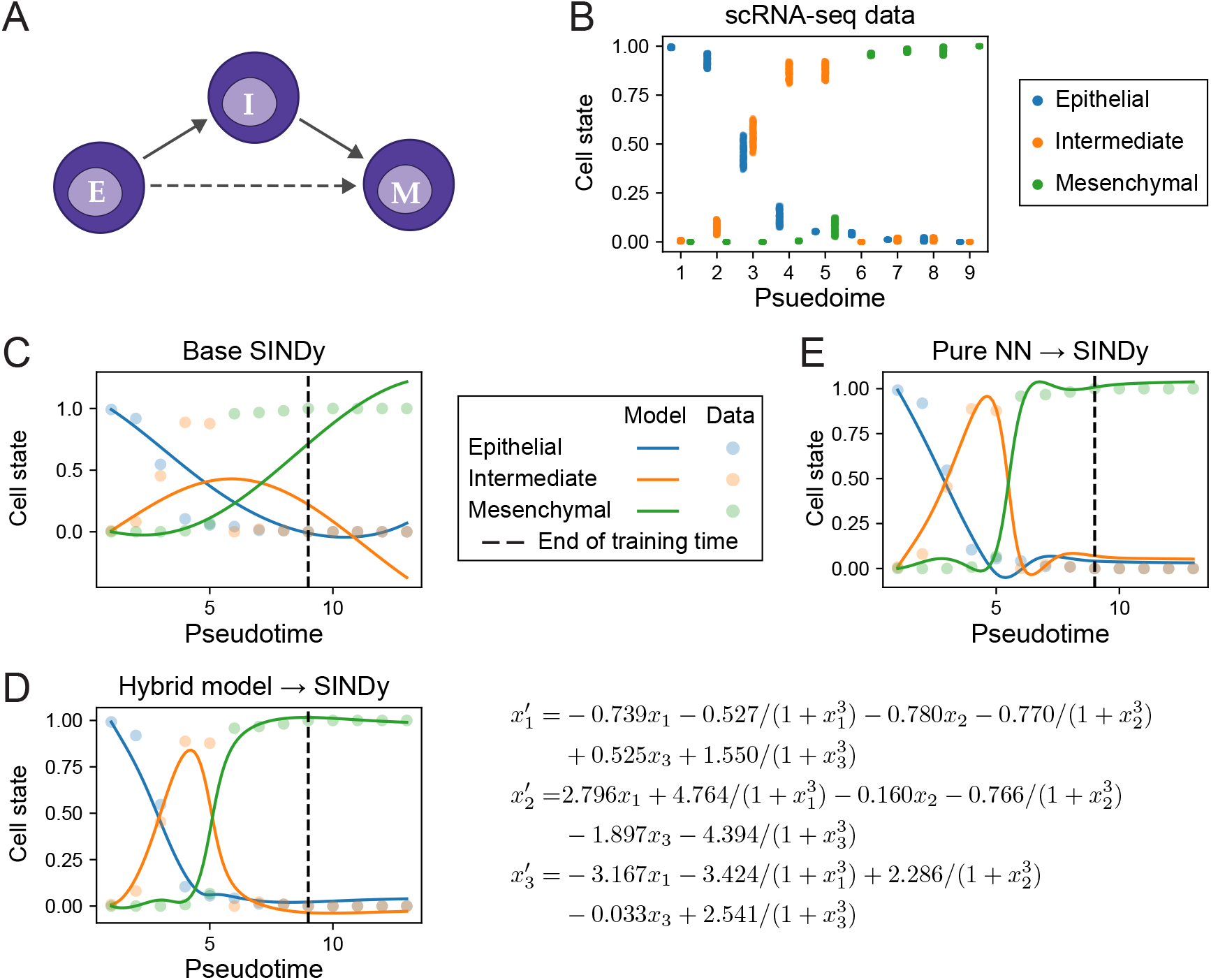
Model discovery of cell state transition dynamics from scRNA-seq data. **A**. Diagram of possible cell state transitions during epithelial-mesenchymal transition (EMT). **B**. EMT data generated with uncertainty estimates for model training from scRNA-seq of cell state transitions over pseudotime. **C**. Simulations from ODE model inferred by base SINDy. **D**. Simulations from ODE model inferred from the hybrid model (Eq 8); model equations of the inferred model (right). *x*_1_: epithelial cell state; *x*_2_ intermediate cell state; *x*_3_: mesenchymal cell state. **E**. Simulations from ODE model inferred from the pure NN model.

To train models, 200 trajectories of the cell population dynamics were sampled over pseudotime values *t* ∈ [1, 9] with step size Δ*t* = 1; for validation of fitted models, 50 additional trajectories were sampled over the same time points. To test inferred models’ ability to extrapolate, 50 trajectories were sampled on *t* ∈ [1, 13] with step size Δ*t* = 1. We first tested base SINDy to directly infer models from the EMT data. We found that it could not produce models with dynamics that fit the data (Fig 7C). This highlights the need for accurate derivatives for SINDy, and the challenges in obtaining these for noisy, sparse data.

We next employed a hybrid dynamical system, assuming that each cell state had a known loss term with a rate parameter of 1, i.e. we fit the model:

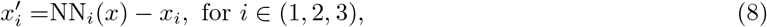

where (*x*_1_, *x*_2_, *x*_3_) denote the epithelial, intermediate, and mesenchymal states, respectively. We inferred models using a two-step approach as above, with the model selection configuration given in S11 Table. The basis libraries consisted of either only polynomial terms or a mixture of polynomial and Hill terms. We compared the models inferred by our hybrid approach with a pure NN approach in which we do not consider any known terms, i.e. we fit the entire derivative by NN(*x*).

Both the hybrid model (Fig 7D) and the pure NN (Fig 7E) could fit the dynamics of EMT. The fits of the hybrid model were slightly better than those obtained by the pure NN model. Analysis of the inferred derivatives by each method identified the critical period in *t* ∈ [3, 6] as the cells transition *E* → *I* and then *I* → *M* (S1 Figure). Comparison of the models inferred from either hybrid (S12 Table) or the pure NN approach (S13 Table) found that the top ranked models from each were consistent, sharing both the same terms topologically and in many cases similar parameter values. The basis of the top inferred model was `hill_3_poly_1:` a mixture of polynomial terms with Hill terms of order 3. Notably, all polynomial terms in the model are linear, indicating a paucity of evidence for higher order interactions. The inferred model contains terms characterizing simple cell state transitions, e.g. *x*_1_ → *x*_2_, albeit with a nonlinear modification to the transition rate for *x*_1_ to *x*_2_ (first two terms of 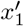 in Fig 7D). The model also suggests that the transition from E to I is also stimulated by the presence of intermediate state cells, and that there is some evidence of a direct reverse transition from M back to E. Finally, it is notable that of the loss terms in Eq 8, the inferred value for the loss rate of *x*_3_ is nearly zero, i.e. SINDy predicted a term of close to 1 (+0.967*x*_3_) canceling out the loss term (−*x*_3_) in the prior. This highlights the ability of the hybrid model framework to “correct” wrongly assumed model terms, given a sufficient basis. Biologically, a loss rate for *x*_3_ of zero seems quite plausible since all cells transition to the mesenchymal state at the end of pseudotime.

## Discussion

In this study, we developed a framework for data-driven discovery of ODE models. We presented methods to infer models from noisy data via a two-step model selection framework. In the first, we learnt the latent (unknown) model dynamics with a NN; in the second we used sparse regression (SINDy) to infer equations to model the system. We showed how the use of hybrid dynamical systems outperformed purely data-driven approaches. This highlights that — while data-driven machine learning approaches for modeling complex systems hold great promise — best performance may be achieved with a degree of supervision when one has some prior knowledge about the system. Analysis of model discovery on two canonical models (Lotka-Volterra and the repressilator) showed that our approach performed well on both additive and multiplicative noise at magnitudes up to 5%. At noise of levels of 10% and above, correct or partially correct models could still be found although not always ranked highest: it can be important to consider a suite of possible models, rather than a single model, and evaluate them comparatively via specific features or interactions. We demonstrated the potential of these methods with application to a canonical cell state transition: where inferred models consisted of relatively simple, primarily first-order cell state transitions.

The primary goal of this study was to identify models with low generalization error; this may come at the expense of underfitting the data, especially in regimes where noise is high. The general question of identifying the underlying dynamics of biological systems with fidelity in the presence of noise remains challenging. Future work might incorporate, for example, inference of stochastic models from the data. There is also room for improvement in inferring models within the current framework, especially for noise levels of 10% and above. One possible way to improve may be to denoise the data using conventional or deep learning-based smoothing techniques for time series [44, 45] before learning a hybrid dynamical model. We also saw that at higher noise levels, incorrect models could be ranked above correct models by AICc. This may be because the AIC term in the AICc grows with the noise, but the correction term does not; future work could explore alternative correction terms to the AIC that explicitly take the noise level into account.

The computational cost of model selection currently grows exponentially with the number of hyperparameters, since a grid search is used to explore the space. This could be reduced by alternative methods for hyperparameter optimization, thus enabling wider explorations of the model space [46, 47]. These algorithms work not only for continuous hyperparameters, but also for discrete or categorical ones, and could be suitable both for NN-based learning or for regression with SINDy. Alternative NN architectures and optimizers for SINDy may also improve future performance, including the SR3 algorithm and Weak SINDy [48–51], and could be incorporated as categorical variables into model selection.

Sources of biological noise are myriad and complex. Yet the most popular algorithms for SINDy have only been tested on zero-mean Gaussian noise [3, 50, 51]. An advantage of this choice is that a simple *L*_2_ norm can be used for the error term in the objective function. However, such an objective function may be unsuitable for data with asymmetric or colored noise. We used the STLSQ optimizer here, considering only datasets with zero-mean Gaussian noise, although it would also be possible to learn the noise empirically. Kaheman et al. [52] proposed a SINDy-based method that fit the noise as a parameter on a per-time point basis, from which it could recover ODE models from measurements with asymmetric noise. However, the number of noise parameters that one must fit is then equal to the number of time points and thus grows with the number of time points in training data. An alternative approach to model the noise that could overcome this issue would be to use (Bayesian) variational inference approaches [53].

Data-driven model discovery combines many fields, from numerical methods and statistical learning to dynamical systems. Its applications are equally broad, spanning engineering and the physical sciences. But applications of model discovery in biology are among the most exciting, precipitated by the astounding quantities of biological data and the extent to which we do not know the structure of most biological networks; there is a lot to learn. Here we have tested the utility of model discovery methods on some challenging problems consisting of data sampled noisily from systems with complex dynamics. We have shown that hybrid methods can successfully learn incomplete models with limited noisy input data, and demonstrated the importance of incorporating prior knowledge into the inference framework. We were also able to predict candidate models describing regulations occurring during EMT from single-cell RNA-sequencing data. We anticipate further development of data-driven model discovery methods to improve our ability to infer models robustly, from limited noisy data, with prior knowledge integration where appropriate. This will accelerate our ability to produce new hypotheses in the form of differential equation models and, eventually, discover new biology.

## Data and code availability

All code associated with this study are available on GitHub at: https://github.com/maclean-lab/model-discovery. Publicly available data (single-cell RNA sequencing of EMT) are available on the gene expression omnibus (GEO) with accession: GSE147405.

## Author contributions

X.W.: Conceptualization, software, investigation, methodology, writing—original draft, and writing— reviewing and editing. M.M.: Investigation, methodology, writing—reviewing and editing. A.L.M.: Conceptualization, investigation, methodology, supervision, writing—original draft, and writing—reviewing and editing.

## Acknowledgements

A.L.M. acknowledges support from the National Institutes of Health (R35GM143019) and the National Science Foundation (DMS 2045327).

