## Supplementary Figures and Tables for "Data-driven model discovery and model selection for noisy biological systems"

Xiaojun Wu[1], MeiLu McDermott[1], Adam L. MacLean[1,*]
[1]Department of Quantitative and Computational Biology, University of Southern California, Los Angeles, CA

### Supplementary Figures

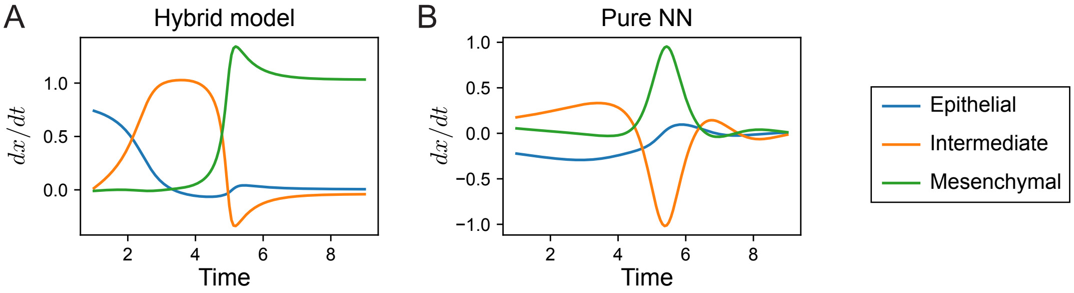


**S1 Figure. Latent dynamics of cell state transition estimated by trained neural networks. A.** Latent dynamics estimated by the trained neural network $\mathrm{NN}(x)$ in hybrid formulation (Eqs 8). **B.** Latent dynamics estimated by the trained neural network $\mathrm{NN}(x)$ in pure neural network formulation (i.e. $x' = \mathrm{NN}(x)$).

### Supplementary Tables

| Noise type | Noise level | Window size | Batch size | Learning rate |
|---|---|---|---|---|
| Additive | 0.1% | 10 | 5 | 0.01 |
| Additive | 0.5% | 5 | 5 | 0.01 |
| Additive | 1% | 5 | 5 | 0.01 |
| Additive | 5% | 5 | 20 | 0.01 |
| Additive | 10% | 10 | 10 | 0.01 |
| Additive | 20% | 5 | 10 | 0.01 |
| Multiplicative | 0.1% | 10 | 10 | 0.1 |
| Multiplicative | 0.5% | 10 | 5 | 0.01 |
| Multiplicative | 1% | 10 | 5 | 0.01 |
| Multiplicative | 5% | 10 | 5 | 0.01 |
| Multiplicative | 10% | 10 | 5 | 0.01 |
| Multiplicative | 20% | 5 | 5 | 0.1 |

**S1 Table. Hyperparameters used to attain lowest mean squared losses for hybrid dynamical models from Lotka-Volterra datasets.**

| Noise type | Noise level | Lowest AICc for given noise model | Inferred model |
|---|---|---|---|
| Additive | 10% | $-145.68$ | $x_1' = 1.3x_1 - 0.841x_1x_2$<br>$x_2' = -1.8x_2 + 0.924x_1x_2$ |
| Additive | 20% | N/A | Not found |
| Multiplicative | 10% | $-58.93$ | $x_1' = 1.3x_1 - 1.063x_1x_2$<br>$x_2' = -1.8x_2 + 0.877x_1x_2$ |
| Multiplicative | 20% | N/A | Not found |

**S2 Table. Best (according to AICc) correct ODE models inferred from Lotka-Volterra datasets using hybrid formulation.** Terms colored in green are known terms in Eqs 4. Models already listed in Table 2 are not shown.

| Noise type | Noise level | Step size | Basis library | $\alpha$ (STLSQ regularization) |
|---|---|---|---|---|
| Additive | 0.1% | 0.1 | `poly_max_2`<br>`poly_2_3` | 0.05<br>0.05, 0.1, 0.5, 1, 5 |
| Additive | 0.5% | 0.1 | `poly_max_2`<br>`poly_2_3` | 0.05, 0.1<br>0.05, 0.1, 0.5, 1, 5 |
| Additive | 1% | 0.05 | `poly_max_2`<br>`poly_2_3` | 0.05, 0.1<br>0.05, 0.1, 0.5 |
| Additive | 5% | 0.05 | `poly_max_2`<br>`poly_2_3` | 0.05, 0.1, 0.5, 1<br>5, 10 |
| Additive | 10% | 0.05 | `poly_2_3` | 0.05, 0.1, 0.5 |
| Additive | 20% | 0.1 | `poly_2_3` | 1 |
| Multiplicative | 0.1% | 0.05 | `poly_2_3` | 0.05, 0.1, 0.5, 1, 5, 10 |
| Multiplicative | 0.5% | 0.1 | `poly_max_2`<br>`poly_2_3` | 0.05, 0.1<br>0.05, 0.1, 0.5, 1, 5 |
| Multiplicative | 1% | 0.1 | `poly_max_2`<br>`poly_2_3` | 0.05, 0.1, 0.5<br>0.05, 0.1, 0.5, 1, 5, 10 |
| Multiplicative | 5% | 0.1 | `poly_max_2` | 0.05, 0.1, 0.5, 1, 5, 10 |
| Multiplicative | 10% | 0.1 | `poly_2_3` | 0.05, 0.1, 0.5 |
| Multiplicative | 20% | 0.1 | `poly_max_2` | 5 |

**S3 Table. Hyperparameters used to attain lowest AICcs for SINDy regression on Lotka-Volterra datasets.**

| Noise type | Noise level | Lowest AICc for given noise model | Inferred model |
|---|---|---|---|
| Additive | 0.1% | $-309.30$ | $x_1' = 1.301x_1 - 0.892x_1x_2$<br>$x_2' = -1.801x_2 + 0.792x_1x_2$ |
| Additive | 0.5% | $-299.04$ | $x_1' = 1.302x_1 - 0.892x_1x_2$<br>$x_2' = -1.800x_2 + 0.790x_1x_2$ |
| Additive | 1% | $-283.38$ | $x_1' = 1.303x_1 - 0.892x_1x_2$<br>$x_2' = -1.799x_2 + 0.786x_1x_2$ |
| Additive | 5% | $-178.24$ | $x_1' = 1.310x_1 - 0.883x_1x_2$<br>$x_2' = -1.782x_2 + 0.728x_1x_2$ |
| Additive | 10% | $-104.66$ | $x_1' = 1.317x_1 - 0.859x_1x_2$<br>$x_2' = -1.759x_2 + 0.653x_1x_2$ |
| Additive | 20% | $-26.46$ | $x_1' = 1.323x_1 - 0.803x_1x_2$<br>$x_2' = -1.732x_2 + 0.585x_1x_2$ |
| Multiplicative | 0.1% | $-297.82$ | $x_1' = 1.301x_1 - 0.892x_1x_2$<br>$x_2' = -1.801x_2 + 0.791x_1x_2$ |
| Multiplicative | 0.5% | $-254.14$ | $x_1' = 1.305x_1 - 0.890x_1x_2$<br>$x_2' = -1.800x_2 + 0.784x_1x_2$ |
| Multiplicative | 1% | $-214.86$ | $x_1' = 1.310x_1 - 0.888x_1x_2$<br>$x_2' = -1.799x_2 + 0.775x_1x_2$ |
| Multiplicative | 5% | N/A | Not found |
| Multiplicative | 10% | N/A | Not found |
| Multiplicative | 20% | N/A | Not found |

**S4 Table. Best (according to AICc) correct ODE models inferred from Lotka-Volterra datasets using base SINDy.**

| Noise type | Noise level | Lowest AICc for given noise model | Inferred model |
|---|---|---|---|
| Additive | 0.1% | −432.58 | $x_1' = 1.297x_1 - 0.909x_1x_2$<br>$x_2' = -1.812x_2 + 0.798x_1x_2$ |
| Additive | 0.5% | N/A | Not found |
| Additive | 1% | −299.50 | $x_1' = 1.274x_1 - 0.869x_1x_2$<br>$x_2' = -1.780x_2 + 0.658x_1x_2$ |
| Additive | 5% | −39.08 | $x_1' = 1.296x_1 - 0.715x_1x_2$<br>$x_2' = -1.772x_2 + 0.746x_1x_2$ |
| Additive | 10% | −147.44 | $x_1' = 1.329x_1 - 0.969x_1x_2$<br>$x_2' = -1.942x_2 + 1.057x_1x_2$ |
| Additive | 20% | N/A | Not found |
| Multiplicative | 0.1% | −332.45 | $x_1' = 1.301x_1 - 0.896x_1x_2$<br>$x_2' = -1.801x_2 + 0.789x_1x_2$ |
| Multiplicative | 0.5% | −318.49 | $x_1' = 1.306x_1 - 0.903x_1x_2$<br>$x_2' = -1.821x_2 + 0.842x_1x_2$ |
| Multiplicative | 1% | −81.38 | $x_1' = 1.310x_1 - 0.817x_1x_2$<br>$x_2' = -1.903x_2 + 0.955x_1x_2$ |
| Multiplicative | 5% | N/A | Not found |
| Multiplicative | 10% | N/A | Not found |
| Multiplicative | 20% | N/A | Not found |

**S5 Table. Best (according to AICc) correct ODE models inferred from Lotka-Volterra datasets using pure neural network formulation.**

| Rank | Hyperparameters | AICc | Inferred model |
|---|---|---|---|
| 1 | Step size: 0.2<br>Basis library: `hill_2`<br>$\alpha$: 10 | $-22.67$ | $x_1' = -2.152/(1+x_2^2) + 9.303/(1+x_3^2)$<br>$x_2' = 7.755/(1+x_1^2)$<br>$x_3' = -1.491/(1+x_1^2) + 10.563/(1+x_2^2) - 1.897/(1+x_3^2)$ |
| 2 | Step size: 0.1<br>Basis library: `hill_1`<br>$\alpha$: 0.05, 0.1, 0.5, 1, 5, 10 | $-20.77$ | $x_1' = -3.861/(1+x_1) - 4.308/(1+x_2) + 13.194/(1+x_3)$<br>$x_2' = 12.594/(1+x_1) - 4.110/(1+x_2) - 3.268/(1+x_3)$<br>$x_3' = -4.412/(1+x_1) + 14.670/(1+x_2) - 5.194/(1+x_3)$ |
| 3 | Step size: 0.2<br>Basis library: `hill_1`<br>$\alpha$: 0.05, 0.1, 0.5, 1, 5, 10 | $-20.67$ | $x_1' = -3.850/(1+x_1) - 4.320/(1+x_2) + 13.193/(1+x_3)$<br>$x_2' = 12.583/(1+x_1) - 4.096/(1+x_2) - 3.277/(1+x_3)$<br>$x_3' = -4.365/(1+x_1) + 14.618/(1+x_2) - 5.186/(1+x_3)$ |
| 4 | Step size: 0.2<br>Basis library: `hill_3`<br>$\alpha$: 0.05, 0.1, 0.5, 1, 5, 10 | $-17.82$ | $x_1' = 9.725/(1+x_3^3)$<br>$x_2' = 9.577/(1+x_1^3)$<br>$x_3' = 10.247/(1+x_2^3)$ |
| 5 | Step size: 0.1<br>Basis library: `hill_3`<br>$\alpha$: 0.05, 0.1, 0.5, 1, 5, 10 | $-17.69$ | $x_1' = 9.724/(1+x_3^3)$<br>$x_2' = 9.615/(1+x_1^3)$<br>$x_3' = 10.253/(1+x_2^3)$ |
| 6 | Step size: 0.1<br>Basis library: `hill_2`<br>$\alpha$: 10 | $-8.60$ | $x_1' = -1.143/(1+x_1^2) - 1.591/(1+x_2^2) + 9.762/(1+x_3^2)$<br>$x_2' = 7.759/(1+x_1^2)$<br>$x_3' = -1.515/(1+x_1^2) + 10.586/(1+x_2^2) - 1.896/(1+x_3^2)$ |
| 7 | Step size: 0.2<br>Basis library: `hill_2`<br>$\alpha$: 5 | $-8.24$ | $x_1' = -1.134/(1+x_1^2) - 1.600/(1+x_2^2) + 9.762/(1+x_3^2)$<br>$x_2' = 7.755/(1+x_1^2)$<br>$x_3' = -1.491/(1+x_1^2) + 10.563/(1+x_2^2) - 1.897/(1+x_3^2)$ |
| 8 | Step size: 0.2<br>Basis library: `hill_2`<br>$\alpha$: 0.05, 0.1, 0.5, 1 | 50.57 | $x_1' = -1.134/(1+x_1^2) - 1.600/(1+x_2^2) + 9.762/(1+x_3^2)$<br>$x_2' = 8.906/(1+x_1^2) - 1.477/(1+x_2^2)$<br>$x_3' = -1.491/(1+x_1^2) + 10.563/(1+x_2^2) - 1.897/(1+x_3^2)$ |
| 9 | Step size: 0.1<br>Basis library: `hill_2`<br>$\alpha$: 0.05, 0.1, 0.5, 1, 5 | 51.22 | $x_1' = -1.143/(1+x_1^2) - 1.591/(1+x_2^2) + 9.762/(1+x_3^2)$<br>$x_2' = 8.926/(1+x_1^2) - 1.487/(1+x_2^2)$<br>$x_3' = -1.515/(1+x_1^2) + 10.586/(1+x_2^2) - 1.896/(1+x_3^2)$ |
| 10 | Step size: 0.1<br>Basis library: `hill_max_3`<br>$\alpha$: 5 | 51.57 | $x_1' = 9.724/(1+x_3^3)$<br>$x_2' = 6.307/(1+x_1) - 1.794/(1+x_1^2) - 2.104/(1+x_3^2)$<br>$+ 5.203/(1+x_1^3) - 1.997/(1+x_2^3)$<br>$x_3' = 1.386/(1+x_2) - 5.322/(1+x_2^2) + 14.700/(1+x_2^3)$ |

**S6 Table. Top 10 (according to AICc) models inferred from repressilator data with 5% additive noise.**

| Rank | Hyperparameters | AICc | Inferred model |
|---|---|---|---|
| 1 | Step size: 0.2<br>Basis library: `hill_2`<br>$\alpha$: 10 | −24.59 | $x_1' = -2.129/(1+x_2^2) + 9.345/(1+x_3^2)$<br>$x_2' = 9.502/(1+x_1^2) - 2.427/(1+x_3^2)$<br>$x_3' = -2.177/(1+x_1^2) + 10.042/(1+x_2^2)$ |
| 2 | Step size: 0.1<br>Basis library: `hill_2`<br>$\alpha$: 0.05, 0.1, 0.5, 1, 5, 10 | −20.58 | $x_1' = -2.123/(1+x_2^2) + 9.340/(1+x_3^2)$<br>$x_2' = 10.004/(1+x_1^2) - 1.203/(1+x_2^2) - 1.877/(1+x_3^2)$<br>$x_3' = -1.692/(1+x_1^2) + 10.513/(1+x_2^2) - 1.152/(1+x_3^2)$ |
| 3 | Step size: 0.2<br>Basis library: `hill_2`<br>$\alpha$: 0.05, 0.1, 0.5, 1, 5 | −20.57 | $x_1' = -2.129/(1+x_2^2) + 9.345/(1+x_3^2)$<br>$x_2' = 9.993/(1+x_1^2) - 1.201/(1+x_2^2) - 1.874/(1+x_3^2)$<br>$x_3' = -1.682/(1+x_1^2) + 10.501/(1+x_2^2) - 1.151/(1+x_3^2)$ |
| 4 | Step size: 0.2<br>Basis library: `hill_1`<br>$\alpha$: 0.05, 0.1, 0.5, 1, 5, 10 | −2.96 | $x_1' = -3.618/(1+x_1) - 4.380/(1+x_2) + 13.044/(1+x_3)$<br>$x_2' = 14.126/(1+x_1) - 4.181/(1+x_2) - 5.105/(1+x_3)$<br>$x_3' = -4.782/(1+x_1) + 14.346/(1+x_2) - 4.176/(1+x_3)$ |
| 5 | Step size: 0.1<br>Basis library: `hill_1`<br>$\alpha$: 0.05, 0.1, 0.5, 1, 5, 10 | −1.96 | $x_1' = -3.621/(1+x_1) - 4.361/(1+x_2) + 13.031/(1+x_3)$<br>$x_2' = 14.139/(1+x_1) - 4.189/(1+x_2) - 5.107/(1+x_3)$<br>$x_3' = -4.805/(1+x_1) + 14.372/(1+x_2) - 4.180/(1+x_3)$ |
| 6 | Step size: 0.2<br>Basis library: `hill_3`<br>$\alpha$: 0.05, 0.1, 0.5, 1, 5, 10 | 28.74 | $x_1' = 10.404/(1+x_3^3)$<br>$x_2' = 9.797/(1+x_1^3)$<br>$x_3' = 10.722/(1+x_2^3)$ |
| 7 | Step size: 0.1<br>Basis library: `hill_3`<br>$\alpha$: | 28.82 | $x_1' = 10.404/(1+x_3^3)$<br>$x_2' = 9.809/(1+x_1^3)$<br>$x_3' = 10.723/(1+x_2^3)$ |
| 8 | Step size: 0.1<br>Basis library: `hill_max_3`<br>$\alpha$: 0.1 | 79.44 | $x_1' = 5.295/(1+x_1) - 4.002/(1+x_1^2) - 5.449/(1+x_2^2)$<br>$+ 3.413/(1+x_2^3) + 9.543/(1+x_3^3)$<br>$x_2' = -34.742/(1+x_1) + 45.179/(1+x_2) + 4.220/(1+x_3)$<br>$+ 45.946/(1+x_1^2) - 49.284/(1+x_2^2) - 16.262/(1+x_3^2)$<br>$- 15.055/(1+x_1^3) + 15.100/(1+x_2^3) + 8.016/(1+x_3^3)$<br>$x_3' = 3.746/(1+x_1) - 6.886/(1+x_1^2) - 2.487/(1+x_3^2)$<br>$+ 3.432/(1+x_1^3) + 10.515/(1+x_2^3) + 2.621/(1+x_3^3)$ |
| 9 | Step size: 0.2<br>Basis library: `hill_max_3`<br>$\alpha$: 0.05 | 79.74 | $x_1' = 5.332/(1+x_1) - 4.026/(1+x_1^2) - 5.516/(1+x_2^2)$<br>$+ 3.466/(1+x_2^3) + 9.541/(1+x_3^3)$<br>$x_2' = -34.252/(1+x_1) + 44.754/(1+x_2) + 4.193/(1+x_3)$<br>$+ 45.436/(1+x_1^2) - 48.847/(1+x_2^2) - 16.224/(1+x_3^2)$<br>$- 14.887/(1+x_1^3) + 14.936/(1+x_2^3) + 7.987/(1+x_3^3)$<br>$x_3' = 3.737/(1+x_1) - 6.872/(1+x_1^2) - 2.497/(1+x_3^2)$<br>$+ 3.429/(1+x_1^3) + 10.516/(1+x_2^3) + 2.633/(1+x_3^3)$ |
| 10 | Step size: 0.2<br>Basis library: `hill_max_3`<br>$\alpha$: 0.1 | 85.98 | $x_1' = 6.907/(1+x_1) - 1.736/(1+x_3) - 4.838/(1+x_1^2)$<br>$- 5.065/(1+x_2^2) + 3.142/(1+x_2^3) + 10.705/(1+x_3^3)$<br>$x_2' = -34.252/(1+x_1) + 44.754/(1+x_2) + 4.193/(1+x_3)$<br>$+ 45.436/(1+x_1^2) - 48.847/(1+x_2^2) - 16.224/(1+x_3^2)$<br>$- 14.887/(1+x_1^3) + 14.936/(1+x_2^3) + 7.987/(1+x_3^3)$<br>$x_3' = 3.737/(1+x_1) - 6.872/(1+x_1^2) - 2.497/(1+x_3^2)$<br>$+ 3.429/(1+x_1^3) + 10.516/(1+x_2^3) + 2.633/(1+x_3^3)$ |

**S7 Table. Top 10 (according to AICc) models inferred from repressilator data with 10% additive noise.**

| Rank | Hyperparameters | AICc | Inferred model |
|---|---|---|---|
| 1 | Step size: 0.2<br>Basis library: `hill_2`<br>$\alpha$: 0.05, 0.1, 0.5, 1 | −25.74 | $x_1' = -1.873/(1+x_2^2) + 9.551/(1+x_3^2)$<br>$x_2' = 10.506/(1+x_1^2) - 1.660/(1+x_2^2) - 1.610/(1+x_3^2)$<br>$x_3' = -1.821/(1+x_1^2) + 10.355/(1+x_2^2) - 1.087/(1+x_3^2)$ |
| 2 | Step size: 0.1<br>Basis library: `hill_2`<br>$\alpha$: 5, 10 | −25.72 | $x_1' = -1.874/(1+x_2^2) + 9.555/(1+x_3^2)$<br>$x_2' = 10.521/(1+x_1^2) - 1.675/(1+x_2^2) - 1.605/(1+x_3^2)$<br>$x_3' = -2.307/(1+x_1^2) + 9.942/(1+x_2^2)$ |
| 3 | Step size: 0.2<br>Basis library: `hill_2`<br>$\alpha$: 5, 10 | −25.71 | $x_1' = -1.873/(1+x_2^2) + 9.551/(1+x_3^2)$<br>$x_2' = 10.506/(1+x_1^2) - 1.660/(1+x_2^2) - 1.610/(1+x_3^2)$<br>$x_3' = -2.290/(1+x_1^2) + 9.923/(1+x_2^2)$ |
| 4 | Step size: 0.1<br>Basis library: `hill_2`<br>$\alpha$: 0.05, 0.1, 0.5, 1 | −25.64 | $x_1' = -1.874/(1+x_2^2) + 9.555/(1+x_3^2)$<br>$x_2' = 10.521/(1+x_1^2) - 1.675/(1+x_2^2) - 1.605/(1+x_3^2)$<br>$x_3' = -1.838/(1+x_1^2) + 10.372/(1+x_2^2) - 1.090/(1+x_3^2)$ |
| 5 | Step size: 0.2<br>Basis library: `hill_1`<br>$\alpha$: 0.05, 0.1, 0.5, 1, 5, 10 | −16.34 | $x_1' = -3.689/(1+x_1) - 4.070/(1+x_2) + 13.008/(1+x_3)$<br>$x_2' = 14.532/(1+x_1) - 4.931/(1+x_2) - 4.623/(1+x_3)$<br>$x_3' = -4.997/(1+x_1) + 14.225/(1+x_2) - 4.097/(1+x_3)$ |
| 6 | Step size: 0.1<br>Basis library: `hill_1`<br>$\alpha$: 0.05, 0.1, 0.5, 1, 5, 10 | −16.13 | $x_1' = -3.678/(1+x_1) - 4.070/(1+x_2) + 13.000/(1+x_3)$<br>$x_2' = 14.541/(1+x_1) - 4.952/(1+x_2) - 4.609/(1+x_3)$<br>$x_3' = -5.034/(1+x_1) + 14.270/(1+x_2) - 4.108/(1+x_3)$ |
| 7 | Step size: 0.2<br>Basis library: `hill_3`<br>$\alpha$: 0.05, 0.1, 0.5, 1, 5, 10 | −0.18 | $x_1' = 11.224/(1+x_3^3)$<br>$x_2' = 10.628/(1+x_1^3)$<br>$x_3' = 10.740/(1+x_2^3)$ |
| 8 | Step size: 0.1<br>Basis library: `hill_3`<br>$\alpha$: 0.05, 0.1, 0.5, 1, 5, 10 | −0.12 | $x_1' = 11.226/(1+x_3^3)$<br>$x_2' = 10.648/(1+x_1^3)$<br>$x_3' = 10.740/(1+x_2^3)$ |
| 9 | Step size: 0.2<br>Basis library: `hill_max_3`<br>$\alpha$: 0.1 | 12.20 | $x_1' = 5.562/(1+x_1) - 2.773/(1+x_3) - 4.082/(1+x_1^3)$<br>$- 1.689/(1+x_2^3) + 11.105/(1+x_3^3)$<br>$x_2' = -58.308/(1+x_1) + 70.190/(1+x_2) - 0.715/(1+x_3)$<br>$+ 64.896/(1+x_1^2) - 76.735/(1+x_2^2) - 3.163/(1+x_3^2)$<br>$- 17.729/(1+x_1^3) + 26.598/(1+x_2^3)$<br>$x_3' = 1.973/(1+x_1) - 2.301/(1+x_1^2) + 10.076/(1+x_2^3)$ |
| 10 | Step size: 0.1<br>Basis library: `hill_max_3`<br>$\alpha$: 0.5 | 58.68 | $x_1' = 5.187/(1+x_1) - 1.217/(1+x_3) - 2.274/(1+x_3^2)$<br>$- 4.063/(1+x_1^3) - 1.861/(1+x_2^3) + 12.083/(1+x_3^3)$<br>$x_2' = -57.836/(1+x_1) + 69.989/(1+x_2) - 0.730/(1+x_3)$<br>$+ 64.231/(1+x_1^2) - 76.587/(1+x_2^2) - 3.195/(1+x_3^2)$<br>$- 17.422/(1+x_1^3) + 26.548/(1+x_2^3)$<br>$x_3' = 10.740/(1+x_2^3)$ |

**S8 Table. Top 10 (according to AICc) models inferred from repressilator data with 10% multiplicative noise.**

| Rank | Hyperparameters | AICc | Inferred model |
|---|---|---|---|
| 1 | Step size: 0.2<br>Basis library: `hill_2`<br>$\alpha$: 10 | −16.97 | $x_1' = 7.800/(1+x_3^2)$<br>$x_2' = 7.514/(1+x_1^2)$<br>$x_3' = 7.401/(1+x_2^2)$ |
| 2 | Step size: 0.2<br>Basis library: `hill_2`<br>$\alpha$: 1, 5 | −1.44 | $x_1' = 7.800/(1+x_3^2)$<br>$x_2' = 10.740/(1+x_1^2) - 1.543/(1+x_2^2) - 1.966/(1+x_3^2)$<br>$x_3' = -1.750/(1+x_1^2) + 8.924/(1+x_2^2)$ |
| 3 | Step size: 0.1<br>Basis library: `hill_2`<br>$\alpha$: 5, 10 | −1.42 | $x_1' = 7.799/(1+x_3^2)$<br>$x_2' = 10.740/(1+x_1^2) - 1.543/(1+x_2^2) - 1.967/(1+x_3^2)$<br>$x_3' = -1.753/(1+x_1^2) + 8.926/(1+x_2^2)$ |
| 4 | Step size: 0.1<br>Basis library: `hill_3`<br>$\alpha$: 0.05, 0.1, 0.5, 1, 5, 10 | 10.77 | $x_1' = 11.973/(1+x_3^3)$<br>$x_2' = 11.165/(1+x_1^3)$<br>$x_3' = 9.406/(1+x_2^3) + 1.521/(1+x_3^3)$ |
| 5 | Step size: 0.2<br>Basis library: `hill_3`<br>$\alpha$: 0.05, 0.1, 0.5, 1, 5 | 10.79 | $x_1' = 11.975/(1+x_3^3)$<br>$x_2' = 11.166/(1+x_1^3)$<br>$x_3' = 9.407/(1+x_2^3) + 1.520/(1+x_3^3)$ |
| 6 | Step size: 0.2<br>Basis library: `hill_max_3`<br>$\alpha$: 10 | 22.41 | $x_1' = 3.012/(1+x_3^2) + 7.410/(1+x_3^3)$<br>$x_2' = 13.640/(1+x_1) - 30.949/(1+x_1^2) + 30.783/(1+x_1^3)$<br>$- 1.220/(1+x_2^3) - 1.957/(1+x_3^3)$<br>$x_3' = 1.476/(1+x_2) + 2.962/(1+x_2^2) - 1.557/(1+x_1^3)$<br>$+ 4.631/(1+x_2^3)$ |
| 7 | Step size: 0.1<br>Basis library: `hill_max_3`<br>$\alpha$: 5 | 22.46 | $x_1' = 3.007/(1+x_3^2) + 7.416/(1+x_3^3)$<br>$x_2' = 13.655/(1+x_1) - 31.008/(1+x_1^2) + 30.838/(1+x_1^3)$<br>$- 1.219/(1+x_2^3) - 1.956/(1+x_3^3)$<br>$x_3' = 1.493/(1+x_2) + 2.920/(1+x_2^2) - 1.560/(1+x_1^3)$<br>$+ 4.659/(1+x_2^3)$ |
| 8 | Step size: 0.1<br>Basis library: `hill_1`<br>$\alpha$: 0.05, 0.1, 0.5, 1, 5, 10 | 39.69 | $x_1' = -4.009/(1+x_1) - 3.588/(1+x_2) + 12.897/(1+x_3)$<br>$x_2' = 14.773/(1+x_1) - 4.593/(1+x_2) - 5.291/(1+x_3)$<br>$x_3' = -4.879/(1+x_1) + 12.309/(1+x_2) - 2.569/(1+x_3)$ |
| 9 | Step size: 0.2<br>Basis library: `hill_1`<br>$\alpha$: 0.05, 0.1, 0.5, 1, 5, 10 | 39.84 | $x_1' = -4.012/(1+x_1) - 3.585/(1+x_2) + 12.896/(1+x_3)$<br>$x_2' = 14.774/(1+x_1) - 4.593/(1+x_2) - 5.291/(1+x_3)$<br>$x_3' = -4.875/(1+x_1) + 12.303/(1+x_2) - 2.568/(1+x_3)$ |
| 10 | Step size: 0.2<br>Basis library: `hill_1`<br>$\alpha$: 0.05, 0.1, 0.5 | 41.12 | $x_1' = -1.620/(1+x_1^2) + 9.418/(1+x_3^2)$<br>$x_2' = 10.740/(1+x_1^2) - 1.543/(1+x_2^2) - 1.966/(1+x_3^2)$<br>$x_3' = -1.750/(1+x_1^2) + 8.924/(1+x_2^2)$ |

**S9 Table. Top 10 (according to AICc) models inferred from repressilator data with 20% additive noise.**

| Rank | Hyperparameters | AICc | Inferred model |
|---|---|---|---|
| 1 | Step size: 0.1<br>Basis library: `hill_2`<br>$\alpha$: 10 | −13.97 | $x_1' = -1.887/(1+x_2^2) + 9.666/(1+x_3^2)$<br>$x_2' = 8.569/(1+x_1^2)$<br>$x_3' = -3.325/(1+x_1^2) + 11.144/(1+x_2^2)$ |
| 2 | Step size: 0.2<br>Basis library: `hill_2`<br>$\alpha$: 5 | −13.95 | $x_1' = -1.884/(1+x_2^2) + 9.657/(1+x_3^2)$<br>$x_2' = 8.564/(1+x_1^2)$<br>$x_3' = -3.311/(1+x_1^2) + 11.124/(1+x_2^2)$ |
| 3 | Step size: 0.2<br>Basis library: `hill_2`<br>$\alpha$: 10 | −13.84 | $x_1' = 7.949/(1+x_3^2)$<br>$x_2' = 8.564/(1+x_1^2)$<br>$x_3' = -3.311/(1+x_1^2) + 11.124/(1+x_2^2)$ |
| 4 | Step size: 0.1<br>Basis library: `hill_2`<br>$\alpha$: 5 | −12.96 | $x_1' = -1.887/(1+x_2^2) + 9.666/(1+x_3^2)$<br>$x_2' = 10.405/(1+x_1^2) - 1.898/(1+x_2^2)$<br>$x_3' = -3.325/(1+x_1^2) + 11.144/(1+x_2^2)$ |
| 5 | Step size: 0.1<br>Basis library: `hill_2`<br>$\alpha$: 0.05, 0.1, 0.5, 1 | −5.81 | $x_1' = -1.887/(1+x_2^2) + 9.666/(1+x_3^2)$<br>$x_2' = 10.948/(1+x_1^2) - 1.333/(1+x_2^2) - 1.132/(1+x_3^2)$<br>$x_3' = -2.764/(1+x_1^2) + 11.728/(1+x_2^2) - 1.168/(1+x_3^2)$ |
| 6 | Step size: 0.2<br>Basis library: `hill_2`<br>$\alpha$: 0.05, 0.1, 0.5, 1 | −5.80 | $x_1' = -1.884/(1+x_2^2) + 9.657/(1+x_3^2)$<br>$x_2' = -1.884/(1+x_2^2) + 9.657/(1+x_3^2)$<br>$x_3' = -2.747/(1+x_1^2) + 11.714/(1+x_2^2) - 1.174/(1+x_3^2)$ |
| 7 | Step size: 0.2<br>Basis library: `hill_3`<br>$\alpha$: 0.05, 0.1, 0.5, 1, 5 | −3.96 | $x_1' = 1.731/(1+x_1^3) + 10.462/(1+x_3^3)$<br>$x_2' = 13.276/(1+x_1^3)$<br>$x_3' = 12.384/(1+x_2^3)$ |
| 8 | Step size: 0.1<br>Basis library: `hill_3`<br>$\alpha$: 0.05, 0.1, 0.5, 1, 5, 10 | −3.47 | $x_1' = 1.712/(1+x_1^3) + 10.492/(1+x_3^3)$<br>$x_2' = 13.280/(1+x_1^3)$<br>$x_3' = 12.394/(1+x_2^3)$ |
| 9 | Step size: 0.2<br>Basis library: `hill_1`<br>$\alpha$: 0.05, 0.1, 0.5, 1, 5, 10 | −1.27 | $x_1' = -2.457/(1+x_1) - 5.191/(1+x_2) + 12.801/(1+x_3)$<br>$x_2' = 14.107/(1+x_1) - 4.479/(1+x_2) - 4.039/(1+x_3)$<br>$x_3' = -6.695/(1+x_1) + 15.906/(1+x_2) - 4.067/(1+x_3)$ |
| 10 | Step size: 0.1<br>Basis library: `hill_1`<br>$\alpha$: 0.05, 0.1, 0.5, 1, 5, 10 | −1.23 | $x_1' = -2.460/(1+x_1) - 5.191/(1+x_2) + 12.807/(1+x_3)$<br>$x_2' = 14.102/(1+x_1) - 4.480/(1+x_2) - 4.032/(1+x_3)$<br>$x_3' = -6.713/(1+x_1) + 15.923/(1+x_2) - 4.065/(1+x_3)$ |

**S10 Table. Top 10 (according to AICc) models inferred from repressilator data with 20% multiplicative noise.**

| Learning step | Hyperparameter | Values/options |
|---|---|---|
| Hybrid dynamical model | Window size (data preprocessing) | 3, 5, 7 |
| | Batch size (data preprocessing) | 5, 10, 20 |
| | Learning rate (Adam optimization) | 0.001, 0.01, 0.1 |
| SINDy | Step size (for generating $\widehat{X}$) | 0.1, 0.25, 0.5, 1 |
| | Basis library | `poly_max_2`, `poly_1_2`, `hill_1_poly_1`, `hill_1_poly_1_xy`, `hill_1_poly_1_2`, `hill_2_poly_1`, `hill_2_poly_1_xy`, `hill_2_poly_1_2`, `hill_3_poly_1`, `hill_3_poly_1_xy`, `hill_3_poly_1_2`, `hill_max_3_poly_1` |
| | $\alpha$ (STLSQ regularization parameter) | 0.05, 0.1, 0.5, 1.0, 5.0, 10.0 |
| | $\lambda$ (STLSQ coefficient threshold) | 1.0 |

**S11 Table. Model selection configuration for EMT data.** For model selection at the SINDy step, we considered basis libraries that include polynomial terms and/or Hill terms — `poly_max_2`: polynomial terms of degree up to 2 (bias term included); `poly_1_2`: polynomial terms of degrees 1 and 2 (no bias term); `hill_n_poly_1`: Hill terms of order $n$ plus linear terms; `hill_n_poly_xy`: Hill terms of order $n$ plus linear terms and interaction terms of the form $xy$; `hill_n_poly_1_2`: Hill terms of order $n$ plus polynomial terms of degrees 1 and 2; and `hill_max_3_poly_1`: Hill terms of order up to 3 plus linear terms.

| Rank | Hyperparameters | AICc | Inferred model |
|---|---|---|---|
| 1 | Step size: 0.1<br>Basis library: `hill_3_poly_1`<br>$\alpha$: 0.05 | −175.35 | $x_1' = -0.527/(1+x_1^3) - 0.770/(1+x_2^3) + 1.550/(1+x_3^3) - 0.739x_1 - 0.780x_2 + 0.525x_3$<br>$x_2' = 4.764/(1+x_1^3) - 0.766/(1+x_2^3) - 4.394/(1+x_3^3) + 2.796x_1 - 0.160x_2 - 1.897x_3$<br>$x_3' = -3.424/(1+x_1^3) + 2.286/(1+x_2^3) + 2.541/(1+x_3^3) - 3.167x_1 - 0.033x_3$ |
| 2 | Step size: 0.5<br>Basis library: `poly_1_2`<br>$\alpha$: 1 | −169.89 | $x_1' = 0.361x_1 - 0.596x_1^2 - 1.742x_1x_2$<br>$x_2' = 1.752x_1 - 6.155x_2 + 37.413x_3 - 1.762x_1^2 + 7.315x_1x_2 + 17.320x_1x_3 + 5.716x_2^2$<br>$- 36.516x_2x_3 - 38.075x_3^2$<br>$x_3' = -0.912x_1 + 9.859x_2 - 32.546x_3 + 0.909x_1^2 - 10.100x_1x_2 - 16.438x_1x_3 - 9.422x_2^2$<br>$+ 26.807x_2x_3 + 33.146x_3^2$ |
| 3 | Step size: 0.25<br>Basis library: `hill_3_poly_1`<br>$\alpha$: 0.05, 0.1 | −160.79 | $x_1' = -0.487/(1+x_1^3) - 0.789/(1+x_2^3) + 1.577/(1+x_3^3) - 0.774x_1 - 0.839x_2 + 0.490x_3$<br>$x_2' = 4.784/(1+x_1^3) - 0.760/(1+x_2^3) - 4.392/(1+x_3^3) + 2.773x_1 - 0.188x_2 - 1.924x_3$<br>$x_3' = -3.427/(1+x_1^3) + 2.247/(1+x_2^3) + 2.566/(1+x_3^3) - 3.146x_1 - 0.004x_3$ |
| 4 | Step size: 0.5<br>Basis library: `poly_1_2`<br>$\alpha$: 0.5 | −140.14 | $x_1' = -0.217x_1 + 0.206x_2 + 2.695x_3 - 1.142x_1x_2 - 0.300x_2^2 - 2.800x_2x_3 - 2.714x_3^2$<br>$x_2' = 1.752x_1 - 6.155x_2 + 37.413x_3 - 1.762x_1^2 + 7.315x_1x_2 + 17.320x_1x_3 + 5.716x_2^2$<br>$- 35.516x_2x_3 - 38.075x_3^2$<br>$x_3' = -0.912x_1 + 9.859x_2 - 32.546x_3 + 0.909x_1^2 - 10.100x_1x_2 - 16.438x_1x_3 - 9.422x_2^2$<br>$+ 26.807x_2x_3 + 33.146x_3^2$ |
| 5 | Step size: 0.1<br>Basis library: `poly_max_2`<br>$\alpha$: 0.5, 1 | −135.98 | $x_1' = 0.559 - 0.808x_1 - 0.974x_2 - 0.549x_1x_2 + 0.338x_2^2 - 0.557x_3^2$<br>$x_2' = -10.299 + 23.287x_1 + 18.400x_2 + 22.126x_3 - 13.028x_1^2 - 18.872x_1x_2 - 10.346x_1x_3$<br>$- 8.286x_2^2 - 23.352x_2x_3 - 12.129x_3^2$<br>$x_3' = 1.364 - 3.568x_1 + 4.043x_2 - 2.735x_3 + 2.219x_1^2 - 3.756x_1x_2 - 7.884x_1x_3 - 5.186x_2^2$<br>$- 0.674x_2x_3 + 1.640x_3^2$ |
| 6 | Step size: 0.5<br>Basis library: `poly_max_2`<br>$\alpha$: 0.1 | −132.74 | $x_1' = 0.528 - 0.774x_1 - 0.922x_2 - 0.572x_1x_2 + 0.316x_2^2 - 0.526x_3^2$<br>$x_2' = -45.538 + 98.131x_1 + 84.636x_2 + 152.511x_3 - 52.612x_1^2 - 88.478x_1x_2 - 71.878x_1x_3$<br>$- 39.635x_2^2 - 150.987x_2x_3 - 107.920x_3^2$<br>$x_3' = 26.971 - 57.993x_1 - 43.913x_2 - 100.714x_3 + 31.025x_1^2 + 46.635x_1x_2 + 36.390x_1x_3$<br>$+ 17.438x_2^2 + 95.196x_2x_3 + 74.512x_3^2$ |
| 7 | Step size: 0.1<br>Basis library: `poly_1_2`<br>$\alpha$: 5, 10 | −132.22 | $x_1' = 0.305x_1 - 0.533x_1^2 - 1.689x_1x_2$<br>$x_2' = 1.342x_1 - 2.027x_2 - 3.614x_3 - 1.363x_1^2 + 2.824x_1x_2 + 9.877x_1x_3 + 1.865x_2^2$<br>$+ 2.369x_2x_3 + 3.375x_3^2$<br>$x_3' = -0.661x_1 + 6.749x_2 + 0.675x_3 + 0.674x_1^2 - 6.630x_1x_2 - 10.563x_1x_3 - 6.531x_2^2$<br>$- 4.081x_2x_3 - 0.414x_3^2$ |
| 8 | Step size: 0.25<br>Basis library: `poly_max_2`<br>$\alpha$: 0.5 | −130.76 | $x_1' = 0.548 - 0.796x_1 - 0.952x_2 - 0.562x_1x_2 + 0.327x_2^2 - 0.546x_3^2$<br>$x_2' = -10.120 + 22.789x_1 + 17.829x_2 + 27.733x_3 - 12.692x_1^2 - 18.079x_1x_2 - 8.776x_1x_3$<br>$- 7.925x_2^2 - 28.893x_2x_3 - 17.975x_3^2$<br>$x_3' = 1.217 - 3.186x_1 + 4.497x_2 - 6.668x_3 + 1.973x_1^2 - 4.357x_1x_2 - 8.976x_1x_3 - 5.471x_2^2$<br>$+ 3.201x_2x_3 + 5.762x_3^2$ |
| 9 | Step size: 0.25<br>Basis library: `hill_3_poly_2`<br>$\alpha$: 0.5 | −121.13 | $x_1' = 0.403/(1+x_3^3) - 0.641x_1 - 0.650x_2 + 0.166x_2^2 - 0.200x_3^2 - 0.715x_1x_2$<br>$x_2' = 2.278/(1+x_1^3) + 10.324/(1+x_2^3) - 9.872/(1+x_3^3) - 3.727x_1 - 7.421x_2 + 11.920x_3$<br>$+ 2.143x_1^2 + 10.151x_2^2 - 19.841x_3^2 + 6.768x_1x_2 + 6.686x_1x_3 - 14.296x_2x_3$<br>$x_3' = 7.403/(1+x_1^3) - 7.980/(1+x_2^3) + 5.362/(1+x_3^3) - 8.804x_1 - 2.034x_3 + 7.744x_1^2$<br>$- 8.928x_2^2 - 1.096x_1x_2 + 6.272x_1x_3 - 3.926x_2x_3$ |
| 10 | Step size: 0.5<br>Basis library: `hill_3_poly_1`<br>$\alpha$: 0.05, 0.1, 0.5, 1 | −120.39 | $x_1' = -0.474/(1+x_1^3) - 0.756/(1+x_2^3) + 1.560/(1+x_3^3) - 0.803x_1 - 0.856x_2 + 0.453x_3$<br>$x_2' = 4.691/(1+x_1^3) - 0.427/(1+x_2^3) - 4.838/(1+x_3^3) + 2.935x_1 + 0.154x_2 - 1.939x_3$<br>$x_3' = -3.381/(1+x_1^3) + 1.962/(1+x_2^3) + 2.900/(1+x_3^3) - 3.210x_1 - 0.210x_2 + 0.065x_3$ |

**S12 Table. Top 10 (according to AICc) models inferred from EMT data using hybrid formulation.**

| Rank | Hyperparameters | AICc | Inferred model |
|---|---|---|---|
| 1 | Step size: 0.1<br>Basis library: `hill_3_poly_1`<br>$\alpha$: 0.05, 0.1 | −220.35 | $x_1' = -0.862/(1+x_2^3) + 1.166/(1+x_3^3) - 0.531x_1 - 0.933x_2 + 0.363x_3$<br>$x_2' = 4.771/(1+x_1^3) - 2.317/(1+x_2^3) - 3.515/(1+x_3^3) + 3.660x_1 - 0.152x_2 - 0.871x_3$<br>$x_3' = -2.259/(1+x_1^3) + 1.066/(1+x_2^3) + 3.105/(1+x_3^3) - 3.023x_1 - 1.246x_2 - 0.105x_3$ |
| 2 | Step size: 0.1<br>Basis library: `hill_1_poly_1`<br>$\alpha$: 0.05 | −212.51 | $x_1' = 0.524/(1+x_1) + 0.973/(1+x_2) - 1.190/(1+x_3) - 0.264x_1 - 0.807x_3$<br>$x_2' = -8.431/(1+x_1) + 4.172/(1+x_2) + 2.788/(1+x_3) - 2.539x_1 + 3.415x_2 + 2.616x_3$<br>$x_3' = 6.036/(1+x_1) - 4.217/(1+x_2) + 1.038/(1+x_3) + 0.180x_1 - 4.679x_2 - 1.978x_3$ |
| 3 | Step size: 0.5<br>Basis library: `hill_1_poly_1_xy`<br>$\alpha$: 0.1 | −178.11 | $x_1' = 0.319/(1+x_1) + 0.229/(1+x_2) - 0.606x_1 - 0.590x_2 - 0.444x_3 - 0.287x_1x_2 - 0.114x_1x_3$<br>$x_2' = -13.567/(1+x_1) + 0.704/(1+x_2) + 15.344/(1+x_3) - 9.080x_1 - 2.087x_2 + 4.739x_3 - 3.102x_1x_2 + 5.558x_1x_3 + 2.289x_2x_3$<br>$x_3' = 3.842/(1+x_1) - 1.042/(1+x_2) - 0.822x_1 - 3.155x_2 - 2.472x_3 + 0.392x_1x_2 - 0.537x_1x_3 + 1.425x_2x_3$ |
| 4 | Step size: 0.25<br>Basis library: `hill_1_poly_1_xy`<br>$\alpha$: 0.05 | −178.09 | $x_1' = 0.158/(1+x_1) - 0.253/(1+x_2) + 0.841/(1+x_3) - 0.885x_1 - 1.010x_2 - 0.223x_3 - 0.509x_1x_2$<br>$x_2' = -13.846/(1+x_1) + 0.767/(1+x_2) + 15.647/(1+x_3) - 9.299x_1 - 2.144x_2 + 4.800x_3 - 3.180x_1x_2 + 5.555x_1x_3 + 2.447x_2x_3$<br>$x_3' = 3.746/(1+x_1) - 0.923/(1+x_2) - 0.890x_1 - 3.116x_2 - 2.498x_3 + 0.376x_1x_2 - 0.445x_1x_3 + 1.455x_2x_3$ |
| 5 | Step size: 0.25<br>Basis library: `hill_2_poly_1_xy`<br>$\alpha$: 0.1 | −176.85 | $x_1' = 0.607/(1+x_1^2) + 0.433/(1+x_2^2) - 0.371/(1+x_3^2) - 0.580x_1 - 0.597x_2 - 0.776x_3 - 0.740x_1x_2 - 0.105x_2x_3$<br>$x_2' = 2.848/(1+x_1^2) - 1.757/(1+x_2^2) - 2.830/(1+x_3^2) + 3.331x_1 + 0.781x_2 + 1.082x_1x_2 + 3.754x_1x_3 - 1.519x_2x_3$<br>$x_3' = 1.399/(1+x_2^2) + 1.374/(1+x_3^2) - 2.709x_1 - 1.933x_2 - 1.778x_3 - 0.920x_1x_2 - 1.867x_1x_3 + 1.308x_2x_3$ |
| 6 | Step size: 0.25<br>Basis library: `hill_1_poly_1_xy`<br>$\alpha$: 1, 5 | −176.39 | $x_1' = 0.118/(1+x_1) - 0.486/(1+x_2) + 1.224/(1+x_3) - 1.016x_1 - 1.225x_2 - 0.145x_3 - 0.610x_1x_2 + 0.175x_1x_3$<br>$x_2' = -14.126/(1+x_1) + 17.053/(1+x_3) - 9.800x_1 - 2.857x_2 + 5.136x_3 - 3.539x_1x_2 + 5.894x_1x_3 + 2.500x_2x_3$<br>$x_3' = 3.746/(1+x_1) - 0.923/(1+x_2) - 0.890x_1 - 3.116x_2 - 2.498x_3 + 0.376x_1x_2 - 0.445x_1x_3 + 1.455x_2x_3$ |
| 7 | Step size: 0.5<br>Basis library: `hill_3_poly_1_xy`<br>$\alpha$: 1, 5 | −174.92 | $x_1' = 0.208/(1+x_3^3) - 0.422x_1 - 0.390x_2 - 0.380x_1x_2 - 0.990x_1x_3$<br>$x_2' = 4.490/(1+x_1^3) - 6.576/(1+x_2^3) + 0.514/(1+x_3^3) + 3.974x_1 - 1.991x_2 + 1.521x_3 + 2.552x_1x_2 + 2.324x_1x_3 + 1.516x_2x_3$<br>$x_3' = -1.674/(1+x_1^3) + 1.344/(1+x_2^3) + 2.252/(1+x_3^3) - 2.700x_1 - 1.108x_2 - 0.484x_3 - 0.400x_1x_2 - 1.694x_1x_3 + 0.138x_2x_3$ |
| 8 | Step size: 0.5<br>Basis library: `hill_3_poly_1_xy`<br>$\alpha$: 0.5 | −173.76 | $x_1' = 0.254/(1+x_3^3) - 0.464x_1 - 0.420x_2 - 0.405x_1x_2 - 1.134x_1x_3 - 0.207x_2x_3$<br>$x_2' = 4.490/(1+x_1^3) - 6.576/(1+x_2^3) + 0.514/(1+x_3^3) + 3.974x_1 - 1.991x_2 + 1.521x_3 + 2.552x_1x_2 + 2.324x_1x_3 + 1.516x_2x_3$<br>$x_3' = -1.674/(1+x_1^3) + 1.344/(1+x_2^3) + 2.252/(1+x_3^3) - 2.700x_1 - 1.108x_2 - 0.484x_3 - 0.400x_1x_2 - 1.694x_1x_3 + 0.138x_2x_3$ |
| 9 | Step size: 0.1<br>Basis library: `hill_2_poly_1_xy`<br>$\alpha$: 0.05 | −170.14 | $x_1' = 0.767/(1+x_1^2) + 0.306/(1+x_2^2) - 0.399/(1+x_3^2) - 0.498x_1 - 0.678x_2 - 0.801x_3 - 0.740x_1x_2$<br>$x_2' = 4.279/(1+x_1^2) - 0.983/(1+x_2^2) - 4.457/(1+x_3^2) + 3.471x_1 + 0.624x_2 - 1.483x_3 + 0.465x_1x_2 + 5.349x_1x_3 - 1.296x_2x_3$<br>$x_3' = 1.380/(1+x_2^2) + 1.386/(1+x_3^2) - 2.700x_1 - 1.937x_2 - 1.764x_3 - 0.919x_1x_2 - 1.901x_1x_3 + 1.311x_2x_3$ |
| 10 | Step size: 0.1<br>Basis library: `hill_1_poly_xy`<br>$\alpha$: 0.5, 1 | −169.56 | $x_1' = 0.291/(1+x_1) + 0.256/(1+x_2) - 0.617x_1 - 0.575x_2 - 0.443x_3 - 0.296x_1x_2 - 0.124x_1x_3$<br>$x_2' = -13.972/(1+x_1) + 0.849/(1+x_2) + 15.707/(1+x_3) - 9.372x_1 - 2.121x_2 + 4.814x_3 - 3.203x_1x_2 + 5.503x_1x_3 + 2.503x_2x_3$<br>$x_3' = 3.475/(1+x_1) - 1.073/(1+x_2) + 0.588/(1+x_3) - 1.190x_1 - 3.351x_2 - 2.378x_3 + 0.227x_1x_2 - 0.305x_1x_3 + 1.565x_2x_3$ |

**S13 Table. Top 10 (according to AICc) models inferred from EMT data using pure neural network formulation.**
